# A Synergistic Desmin-SPARC Axis Influences Cardiac Stem Cell Differentiation and Promotes Cardiomyogenesis through Autocrine Regulation

**DOI:** 10.1101/2024.03.28.587296

**Authors:** Lucia Leitner, Martina Schultheis, Franziska Hofstetter, Claudia Rudolf, Valeria Kizner, Kerstin Fiedler, Marie-Therese Konrad, Julia Höbaus, Marco Genini, Julia Kober, Elisabeth Ableitner, Teresa Gmaschitz, Diana Walder, Georg Weitzer

## Abstract

**BACKGROUND:** The mammalian heart contains cardiac stem cells throughout life, but it has not been possible to harness or stimulate these cells to repair damaged myocardium in vivo. Assuming physiological relevance of these cells, which have evolved and have been maintained throughout evolution, we are investigating their function using mouse cardiac stem cell lines as an in vitro model system.

**METHODS:** Here we use genetically modified embryonic stem cells and cardiac stem cells from the mouse as model systems to study the influence of desmin and Secreted Protein Acidic and Rich in Cysteine (SPARC) on cardiomyogenesis in embryoid bodies and cardiac bodies. We analyze their expression in self-renewing and differentiating stem cells by fluorescence microscopy, RT-qPCR, quantitative Western blotting and fluorescence activated cell sorting, and assess their influence on the expression of myocardial transcription factors.

**RESULTS:** In embryoid bodies, desmin induces expression and secretion of SPARC, which promotes cardiomyogenesis. Cardiac stem cells secrete substantial amounts of SPARC, which also promotes cardiomyogenesis in a concentration-dependent, autocrine manner and promotes expression of myocardial transcription factors and *desmin*. Desmin and SPARC interact genetically and form a positive feedback loop and secreted SPARC negatively influences sparc mRNA expression. Finally, SPARC rescues cardiomyogenic desmin-haploinsufficiency in cardiac stem cells in a glycosylation-dependent manner, increases the phosphorylation of Smad2 and induces the expression of *gata4, nkx2.5* and *mef2C*.

**CONCLUSIONS:** Demonstration that desmin-induced autocrine secretion of SPARC in cardiac stem cells promotes cardiomyogenesis raises the possibility that a physiological function of cardiac stem cells in the adult and aging heart may be the gland-like secretion of factors such as SPARC that modulate age-related and adverse environmental influences and thereby contribute to cardiac homeostasis throughout life.

## Introduction

Disruption of the network regulating gene expression during cardiogenesis in mammals, including humans, causes various congenital heart defects ^1^. The core unit of highly conserved transcription factor genes in this gene regulatory network, named cardiac kernel ^2^, has been reconstructed, confirmed and expanded from enhancer occupancy studies ^3^. Mutations in genes of the cardiac kernel, such as *gata4*, *tbx5* or *nkx2.5*, cause a range of CHDs with comparable pathologies ^4–6^. However, the highly pleiotropic pathologies associated with mutations within these loci, suggest that these genes are modulated by a variety of developmental stage and homeostatic condition-dependent co-factors. Among the plethora of growth factors influencing cardiomyogenesis and the expression of the cardiac kernel genes in developmental stage-dependent manners two other candidate co-factors emerged. These factors are the muscle-specific intermediate filament protein desmin ^7^ and the matri-cellular protein Secreted Protein Acidic and Rich in Cysteine (SPARC) ^8^. Both proteins are expressed very early during mouse embryogenesis in cardiac progenitor cells ^9–11^ and in adult mouse heart-derived cardiac stem cells (CSCs) in a developmental stage-dependent and cell type-specific fashion ^12, 13^. They both also influence *nkx2.5* expression and cardiomyogenesis in embryonic stem cell (ESC)-derived embryoid bodies (EBs) ^14–16^.

The main and well-established function of desmin is to serve as a subunit for the assembly of the type III intermediate filament network in muscle cells, which in turn functionally connects the z-disks of sarcomeres in myofibrils with mitochondria, the outer nuclear membrane, and with costameres and intercalated disks of the myocardial sarcolemma ^7, 17^. Desmin further promotes cardiomyogenesis in ESC-derived cardiac progenitor cells in EBs and positively influences the expression of *nkx2.5* and *brachyury* ^15, 18^. In CSCs, desmin influences the expression of the *nkx2.5* gene by migration to the nucleus and physical interaction with its minimal cardiac-specific enhancer and the proximal promoter region. Overexpression of *desmin* rescues the nkx2.5-haploinsufficiency phenotype in cardiac progenitor cells in EBs ^16^. Vice versa, overexpression of *nkx2.5* increases the expression of *desmin* in mesenchymal stem cell-derived cardiac progenitor cells ^19^. *Desmin* knockout in mice leads to fibrosis, calcification, and mitochondrial malformation in the adult heart and to sudden death due to rupture of the myocardium under stress conditions ^20–22^. In humans, mutations in the *desmin* genes cause a broad spectrum of myocardial defects leading to end-stage heart diseases which are collectively referred to as desminopathies ^17, 23, 24^.

SPARC is a typical matri-cellular protein influencing extracellular matrix (ECM) composition, cell adhesion and migration, Tgf-ß signaling, and bone mineralization ^25^. It is expressed in a variety of cell types and tissues ^26^. In particular it is expressed by cardiac fibroblasts and seems to be involved in cardiac fibrosis, inflammation, and the CSC niche establishment ^8, 27, 28^. SPARC expression significantly contributes to age-related cardiac dysfunction in *Drosophila melanogaster* ^29, 30^ and mice ^28^, and was downregulated in a patient with dilated cardiomyopathy ^31^. Knockout of *sparc* in mice also results in increased cardiac rupture and dysfunction after acute myocardial infarction ^32^. In humans, evidence emerges that SPARC expression contributes to the etiology and progression of dilated cardiomyopathies ^33^. Similar to desmin, parietal endoderm-secreted SPARC activates *nkx2.5* expression in ESC-derived cardiac progenitor cells and in primary cardiomyocytes from newborn mice and promotes cardiomyogenesis in EBs ^14^. Intriguingly, stress-induced cardiac pathology in aged desmin-null mice ^21^ and in SPARC-null mice after induced myocardial infarction ^32^ is very similar and may hint at a synergistic or overlapping role of both proteins in the adult and aging heart. We therefore sought to investigate the molecular mechanisms by which SPARC and desmin contribute to cardiomyogenesis in CSCs.

Here we show that desmin promotes cardiomyogenesis also in a non-cell autonomous manner by enhancing the expression and secretion of SPARC. SPARC in turn promotes cardiomyogenesis in EBs in a Tgf-ß signal pathway-dependent manner and in CSC-derived cardiac bodies (CBs), where it influences *desmin* expression in a concertation-dependent manner. Crosswise genetic interactions of the *desmin* and *sparc* genes establish a positive feedback loop in CSCs, which, together with a paracrine SPARC-dependent negative feedback loop, regulates cardiomyogenesis in differentiating CSCs in vitro. Knockout of one allele of *sparc* or *desmin* significantly delays and diminishes cardiomyogenesis and causes a very similar cardiomyogenic haplo-insufficient phenotype in CBs, which can be partially rescued by paracrine glycosylated SPARC in SPARC-deficient CBs and completely rescued in desmin-deficient CBs. Finally, paracrine and autocrine SPARC promotes the phosphorylation of Smad2 and the expression of cardiomyogenic transcription factor genes *gata4*, *nkx2.5*, and *mef2C*. These molecular data together with the fact that adult desmin-and SPARC-null mice exhibit similar stress-induced cardiac phenotypes suggest that this desmin-SPARC interplay at the molecular level may support homeostasis or myocardial regeneration in the adult, diseased, and aging heart in vivo.

## Materials and Methods

### Ethics Statement

Hearts for histological analysis and cardiomyocyte isolation were obtained from mice that were humanely maintained and sacrificed in accordance with the Austrian Animal Welfare Act (Bundesgesetz über den Schutz der Tiere (Tierschutzgesetz - TSchG), Art.2 §6 Abs.3, BGBl. I Nr. 118/2004 zuletzt geändert durch BGBl. I Nr. 2/2024). According to this legislation no ethical review or approval is required for the breeding and sacrificing of mice at academic institutions such as the Medical University of Vienna.

### Cell culture

SNL76/7 fibroblasts ^34^ or STO cells ^35^ were maintained in DMEM supplemented with 2mmol/L glutamine, 0.05mg/ml streptomycin, 0.03mg/ml penicillin and 10% (v/v) fetal bovine serum (FBS) (Gibco 26140079, Thermo Fisher Scientific) (M10). Murine AB2.2 and W4 ESCs ^36^ and A5 and He2 cardiac stem cells (CSCs, formerly CVPCs for cardiovascular progenitor cells) ^12^ and all genetically modified derivatives of these ESC and CSC lines (generation procedure described below) were cultured on SNL76/7 feeder cells in DMEM supplemented with 2mmol/L glutamine, 0.05mg/ml streptomycin, 0.03mg/ml penicillin, 0.1mmol/l ß-mercaptoethanol and 15% (v/v) FBS (HyClone HYCLSH30088.03, VWR and SH30071.03, Cytiva) (M15Hy). Feeder cells were generated from SNL76/7, STO or CSCs by incubation with 10μg/ml Mitomycin C for 4 hours.

### Generation of genetically modified ESC and CSC lines

AB2.2 ESC lines with the mutant *desmin* allele *des^-/-^* or *des^+/+/^D^ect^*^15, 37^, the parental A5 wild-type ^12^ and A5 CSC lines expressing *desmin-mCherry*, *mCherry-desmin* or *mCherry* ^16^ have been previously generated and characterized. Here, two *sparc^+/-^*lines (A5 K11 and A5 K17), one *sparc^+/-^* line (He2 K2), as well as four *des^+/-^* lines (A5 1D, 2H, 1E and 3B) CSCs were generated by homologous recombination with a *desmin* knockout vector ^37^ and a *sparc* knockout vector ^38^, respectively, as previously described ^16, 39^. A5 *sparc^+/+^SmC^ect^* (K5 and K6), A5 *sparc^+/+^mC^ect^*(K28 and K46), A5 *sparc^+/+^S^ect^* (K4 and K10), STO *sparc^ect^*, SNL76/7 *sparc^ect^*, 3T3 *sparc^ect^*, and A5 *des^+/+^D^ect^* (K4 and K12) ^16^ were generated by non-homologous recombination with a pCMV3-SPARC *sparc* expression cassette (SinoBiological, MG50494-UT) and a *desmin* expression cassette ^15^, respectively, as previously described ^15, 16, 39^. A5 CSC and W4 ESC reporter cell lines for *brachyury* and *mesp1* promoter activation were generated by non-homologous recombination with mouse 1.2kb pT-Bra^P^-Puro-IRES2-eGFP plasmid ^40^ and human 3.4 kb pMesP1-eGFP ^41^, respectively. A5 and W4 *nkx2.5* reporter cell lines were generated by homologous recombination with a knock-in plasmid pCSX-EGFP-PPDT ^42^ as previously described ^16^. All cell lines were tested for correct expression of the transgenes.

Plasmid DNA was isolated and purified from XL1-blue *E. coli* using QuiaShredders (Quiagen, 79656) and a Quiagen Minipre Kit (Quiagen, 12123). 1.1x107 cells were electroporated at 230V using 500μF with a time constant of 5.6 to 7.0 milliseconds; 25μg linearized DNA and positive clones were selected with G418, puromycin or hygromycin.

### Analysis of cardiomyogenesis in EBs and CBs

Differentiation of ESCs and CSCs was achieved by aggregation of 800 cells to embryoid bodies (EBs) and cardiac bodies (CBs) in hanging drop cultures in DMEM supplemented with 2mmol/L glutamine, 0.05mg/ml streptomycin, 0.03mg/ml penicillin, and 15% (v/v) FBS (Sigma F7524, Sigma-Aldrich) (M15Si) as previously described ^12, 15, 16^. Briefly, ESCs or CSCs were grown to confluence on feeder cells and split 1:2 on the day before aggregation. The next day, cells were dissociated with trypsin and suspended in M15Si medium. Feeder cells were removed by adsorption on gelatin-coated tissue culture plates at 37°C for 45 minutes ^43^. Cells remaining in the supernatant were diluted to 4.5x10^4^ cells/ml M15Si and 90 or 33 20μl drops were placed on the lids of bacterial grade 10cm or 6cm plates. Hanging drops were incubated over a layer of sterile water to compensate for the increased vapor pressure of the drops at 37°C for 4.5 (EBs) or 4.8 (CBs) days. Aggregates were then washed from the lid into gelatin-coated 10cm or 6cm tissue culture plates with 6ml or 4ml of M15Si, evenly distributed, and cultured using a feeding protocol exactly as described in ^14, 43^. From day 16 onwards, medium was changed daily. EBs and CBs were cultured in the presence of either 3mg/l recombinant SPARC (S5174-25UG, Sigma-Aldrich; 942-SP-050, R&D Systems), various amounts of anti-SPARC antibodies (Santa Cruz, (H-90): sc-25574 Sigma, for EBs; PA5-89864, Thermo Fischer Scientific, for CBs), conditioned media from STO cells, various CSC lines, EBs or CBs (see EB and CB co-cultures and media conditioning), 3.5µmol/l SB431242 (S4317, Sigma-Aldrich), or AcSDKP (350461, Abbiotec) for different time periods, as indicated in the figure legends.

Cardiomyogenesis was assessed by calculating the percentage of EBs or CBs with one or more clusters of spontaneously contracting cardiomyocytes daily from day 10 to a maximum of day 65. This provided information on the onset of cardiomyogenesis, the rate of progression, the peak with the maximum percentage of contracting EBs or CBs, and the decline, persistence or longevity of cardiomyogenesis. Counting the total number of contracting clusters from 5 days before to 5 days after peak contraction activity allowed assessment of the extent of the commitment to cardiomyogenic lineages and determination of contraction frequency. The percentage of arrhythmically contracting cells over the same period was also used to assess the quality of the developing cardiomyocytes.

### EB and CB co-cultures and conditioning of media

To co-culture different genotypes of EBs or CBs, respectively, they were transferred from the hanging drop culture to either the well or the sieve insert of 24-well or 6-well tissue culture plates. Both the wells and the sieve inserts had been coated with a 0.1% gelatin solution two hours before.

Different types of conditioned media were prepared to be used for co-culture and to determine the concentration of secreted SPARC. STO and 3T3 cells were cultured at a density of 70,000 cells per cm^2^ and ml of medium, ESC or CSC cells at a density of 500,000 cells per well of 24-well plates, and EBs or CBs at a density of 0.57 CBs/EBs per cm^2^ and 0.35ml of medium. Conditioning was always performed for 24 or 48 hours.

### Quantification of SPARC in media

Absolute levels of SPARC in conditioned media were determined by immune-dot blot assays. EBs were cultured in M15Hy medium for 48 hours and serial dilutions of this medium were spotted onto a nitrocellulose membrane. The membrane was blocked with 5% milk powder in TBS containing 0.1% Tween for 45 minutes. SPARC was detected with rabbit anti-SPARC antibody (#sc-25574, Santa Cruz, 1:200) and secondary alkaline phosphatase-conjugated anti-rabbit IgG antibody (#111-035-003, Dianova, 1:5000). Serial dilutions from 2μg to 0.0625μg of recombinant SPARC (SRP3159, Sigma-Aldrich; S5174-25UG, Sigma-Aldrich; 942-SP-050, R&D Systems) were spotted and used as standards. The relative amounts of SPARC secreted by different genotypes of CSCs were determined by quantitative Western blotting as described below.

### Isolation, detection and quantification of proteins

CSCs were cultured either on feeder cells or on gelatin-coated tissue culture plates for 24 hours after feeder cells were removed by adsorption on gelatin-coated tissue culture plates at 37°C for 45 minutes ^43^. CSCs or cells from CBs were dissociated by incubation with trypsin for 20 minutes, re-suspended in M15Hy, counted on a CellDrop FL (DeNovix), washed once in 1ml PBS and lysed in lysis buffer containing 20mM Tris, pH 7.4, 150mM NaCl, 500mM EDTA, 0.1% NP40, 10% glycerol, 1% EDTA-free protease inhibitor cocktail (B14001, Bimake) and 1% phosphatase inhibitor cocktail (B15001, Bimake) at 4°C for 20 minutes. Insoluble material was removed by centrifugation at 14,000rpm for 20 minutes at 4°C and the supernatants were stored at -80°C. The volume of lysis buffer was adjusted to the total cell number, resulting in a constant concentration of 6000 cell equivalents of protein in the lysate. Protein samples were mixed with 4X Protein Sample Loading Buffer (928-40004, LiCOR), boiled at 95°C for 5 minutes, centrifuged at 14,000rpm at RT for 2 minutes and separated on precast 10% SDS-PAGE gels (4561036, Bio-Rad) at 100V for 70 minutes. Proteins were transferred to nitrocellulose blotting membranes (10-6000-01, VWR) in Harlow buffer at 350mV and 4°C for 2 hours. The membrane was then washed in TBS and stained with Revert™ 700 Total Protein Stain and Wash Solution (926-11015, LiCOR) and analyzed using an Odyssey CLx Fluorescence Imager (LiCOR).

For immune-detection, membranes were washed in TBS, then blocked with 5% milk powder in TBS at RT for 1 hour and incubated with the primary antibody (**Major Resource Table**) in 5% milk powder in TBS at 4°C over night. After washing off of the primary antibodies 3 times with TBST for 10 minutes each, membranes were incubated with the secondary antibody (**Major Resource Table**) in 5% milk powder in TBST at 4°C in the dark for 1 hour, then washed 3 times in the dark with TBST for 10 minutes each and analyzed with an Odyssey CLx Fluorescence Imager (LiCOR). Proteins were quantified in the histogram mode of Photoshop CS2 and relative amounts of proteins were calculated by multiplication of the band size in pixels and the mean luminosity of the band with Microsoft Excel 2013.

### RNA isolation and cDNA synthesis

Total RNA was extracted from single cell suspensions of 500,000 CSCs or CB-derived cells using the FavorPrep™ Blood/Cultured Cell Total RNA Mini Kit (Favorgen, FABRK001-2), eluted with 40μl RNase-free ddH2O and the RNA concentration was measured with a Nanodrop 2000c spectrophotometer (Thermo Scientific). RNA samples were stored at -70°C. cDNA was prepared from 1µg of RNA using the LunaScript® RT SuperMix Kit (New England Biolabs, E3010L). A no template control (NTC) and no reverse transcriptase (NRT) control were performed. Samples were diluted 1:50 and stored at -20°C. mRNA from ESCs was isolated using the Quiagen RNeasy Kit (Quiagen, 74104) and cDNA was synthesized using the RevertAid First Strand cDNA Synthesis Kit (K1622, Thermo Fisher Scientific).

### PCR and qPCR

PCR was performed using Platinum™ SuperFi II Green PCR Master Mix (12369010, Thermo Fisher Scientific) and also with PCR Master Mix (M7502, Promega), Taq polymerase (EP0402, Thermo Scientific) or Pyrococcus furiosus polymerase (EP0501, Thermo Scientific), and 10mM dNTP mix (71005M, Fermentas, Sigma-Aldrich,) with 0.5µl of 10µM primer mix (**Major Resource Table**) in a volume of 25µl. The number of cycles for each primer pair was carefully determined by several preliminary experiments and chosen so that none of the resulting signals were saturated. Samples were run on 1.5% to 2.5% agarose gels and DNA was stained with ethidium bromide.

qPCR was performed for 3 biological replicates with 2 technical replicates each, using the Luna® Universal qPCR Master Mix Protocol (M3003E, New England Biolabs) in 96-well Biorad qPCR plates and a CFX384 Touch Real-Time PCR Detection System (Biorad) or a CFX Opus 96 Real-Time PCR System (Biorad). Each reaction contained 10µl Luna® Master Mix, 4µl H2O, 1µl of a 10µM primer mix, and 5µl of the 1:50 diluted cDNA. The qPCR cycling protocol was 95°C for 60 seconds, then 39 cycles of 95°C for 15 seconds plus 60°C for 30 seconds and finally a step gradient of 0.5°C per 5 seconds from 65 to 95°C before cooling to 4°C. Raw Cq values were exported and expression levels were calculated using the delta-Cq method, with *RPL32* as a housekeeping gene for normalization, and statistical analysis was performed in Microsoft Excel 2013.

### Fluorescence-activated cell sorting

ESCs and CSCs were cultured in triplicate on SNL76/7 feeder cells in M15Hy on 24-well plates for 24 hours, then the medium was replaced with conditioned medium from confluent SPARC-expressing STO or 3T3 cells, and wild-type STO or 3T3 cells as control, for 6 hours and 24 hours, respectively. After incubation with trypsin solution (27250018, Invitrogen) for 20 minutes at 37°C, cells were resuspended in 1ml M15Hy and the number of EGFP-positive cells was analyzed on a BD FACSCalibur™ flow cytometer using BD CellQuest™ Pro software. Data from all three measurements per plate were converted using the FlowJo v10.4 platform and exported to Microsoft Excel 2013 for manual gating and statistical analysis.

### Reporter-gene assays

CSCs and cells from 9-day-old CBs were transiently transfected with the *nkx2.5* promoter-reporter plasmid pNKE24 ^44^ and with pGL3-Basic (E1751, Promega) and phRL-TK (E2241, Promega) as controls and analyzed for promoter activation in the presence of 3mg/l recombinant human SPARC (SRP3159, Sigma-Aldrich) using the Dual-Luciferase® Reporter Assay System (E1910, Promega) and the ONE-Glo™ Luciferase Assay System (E6110, Promega) exactly as described in ^16^.

### Confocal immunofluorescence microscopy and fluorimetry

CSCs were processed and stained as described previously ^15^. Briefly, cells were fixed in 4% paraformaldehyde in PBS for 20 minutes at room temperature, permeabilized with 0.15% saponin in PBS, and stained with antibodies against SPARC (#sc-25574, Santa Cruz, 1:500) for 60 minutes. They were then washed three times with PBS and stained with Alexa488-conjugated secondary antibody (A-11001, Molecular Probes, 1:1000) for 60 minutes. The nuclear DNA was then stained with DAPI (D9542, Sigma) for 5 minutes.

For immunolocalization of SPARC-positive cells in mouse hearts, mice were killed by cervical dislocation. Hearts were dissected, rinsed with PBS, and fixed in 4% formaldehyde in PBS at 4°C over night. Fixed hearts were again rinsed with PBS and placed in an Excelsior Tissue Processor (Histocom) for standard dehydration with ethanol and xylol for a total of 9 hours and then placed in a Histostar embedding station (Histocom) for paraffin embedding for 4 hours. Hearts were sectioned on an RM2255 microtome (Leica) at 3μm section thickness, paraffin embedded, and mounted on glass slides. Heart sections were blocked with 5% BSA in PBS for 1 hour at room temperature. Slides were then incubated for 1 hour with rabbit anti-SPARC primary antibody (PA5-89864, Thermo Fischer Scientific, 1:100) and Alexa Fluor 488 anti-mouse Ly-6A/E (Sca-1) antibody (108115, BioLegend, 1:1000) diluted in PBS, washed three times with PBS, then incubated with secondary IRDye 800CW goat anti-rabbit IgG antibody (926-32211, LICOR, 1:800) for 1 hour and finally washed three times with PBS. Sections were mounted in Fluoromount G mounting medium containing DAPI (00-4958-02, Thermo Fischer Scientific) and imaged using a Zeiss Axio Imager Z2. For visualization of the SPARC-mCherry or mCherry-desmin protein, CSCs and their differentiating derivatives were fixed with absolute ethanol at -20°C for 5 minutes. Photomicrographs were taken on a Zeiss LSM 510 confocal microscope at 615nm.

For quantification of SPARC-mCherry protein internalized by cells and ECM-bound mCherry protein, photomicrographs were taken using a Zeiss Axivert 135 TV microscope, and mCherry fluorescence was quantified using Adobe Photoshop software. To quantify mCherry fluorescence, images obtained with a 615nm filter were imported into Photoshop CS2 as raw, unsaturated 16-bit images, and pixel intensity values within different areas of the ECM, cells, and nuclei, respectively, were extracted using the built-in histogram tool. Pixel intensities were subtracted from the background pixel intensity obtained from multiple cell-or nucleus-free areas of the images.

SPARC-mCherry fluorescence in the medium was measured using a fluorimeter at 610nm with an excitation wavelength of 590nm.

### Statistical analysis

All data are presented as the arithmetic mean ± standard deviation σ_x(n-1)_. Statistical significance was evaluated using Student’s t-test and values of p ≤ 0.05 were considered to indicate statistical significance.

## Results

### Desmin-induced SPARC expression promotes cardiomyogenesis in embryoid bodies

In the past we have shown that desmin influences in vitro cardiomyogenesis in a cell-autonomous and expression level-dependent manner ^15, 16^. To test the hypothesis that desmin also influences cardiomyogenesis in a non-cell-autonomous manner we designed a co-culture experiment in which *des^+/+^D^ect^* ESC-derived EBs were grown on sieve inserts, while either wild-type *des^+/+^* EBs or *des^-/-^* EBs were grown underneath on a separate layer, allowing the desmin-overexpressing cells to act on them in a paracrine manner (**Figure 1A**). Cardiomyogenesis was monitored daily and quantified by calculating the percentage of EBs with rhythmically contracting clusters of cardiomyocytes. We demonstrated that desmin-overexpression in *des^+/+^D^ect^*ESC-derived EBs promotes cardiomyogenesis in co-cultured wild-type *des^+/+^*EBs and partially rescues cardiomyogenesis in *des^-/-^* EBs in a paracrine manner. We identified secreted protein acidic and rich in cysteine (SPARC) as potential candidate mediating the effect of desmin on cardiomyogenesis in a paracrine manner by testing *des^+/+^D^ect^*ESC-derived EBs for increased expression of secreted factors using RT-sqPCR (**Figure 1B**). Paracrine SPARC has previously been shown to promote cardiomyogenesis in EBs ^14^, and quantification of SPARC protein secreted into the medium of *des^+/+^D^ect^*EBs or from isolated parietal endoderm cells showed that *desmin* overexpression also caused increased secretion of SPARC, whereas SPARC secretion was reduced in *des^-/-^* EBs (**Figure 1C**). To demonstrate that it is SPARC that promotes cardiomyogenesis in wild-type *des^+/+^* and in *des^-/-^* EBs, we repeated the co-culture experiment in the presence of SPARC-neutralizing quantities of polyclonal anti-SPARC antibodies (**Figure 1D**). Anti-SPARC antibodies partially inhibited cardiomyogenesis in *des^+/+^*EBs when co-cultured with either *des^+/+^* or *des^+/+^D^ect^*EBs, and strongly inhibited cardiomyogenesis induced by SPARC-secreting *des^+/+^D^ect^*EBs in *des^-/-^* EBs. Because SPARC has been shown to induce Smad2 phosphorylation in the myocardium ^28^, we tested whether anti-SPARC antibodies are able to counteract the paracrine SPARC-mediated increase of phosphorylated Smad2 in *des^-/-^* EBs (**Figure 1E**). Indeed, SPARC increased the phosphorylation of Smad2 in *des^-/-^* EBs, whereas addition of anti-SPARC antibodies partially inhibited Smad2 phosphorylation. Simultaneous addition of recombinant SPARC and SB431542, an inhibitor of Tgf-ß receptor ALK5, also abolished the positive effect of paracrine SPARC on cardiomyogenesis (**Figure 1F**), and inhibition of nuclear import of pSmad2 by AcSDKP in *des^+/+^* EBs also partially inhibited cardiomyogenesis (**Figure 1G**). From these data we may conclude that increased *desmin* expression in differentiating ESCs induces *sparc* expression and that SPARC contributes to cardiomyogenesis in a paracrine and partially ALK5 and Smad2 dependent manner. The major limitation of these experiments in EBs is that we cannot obtain any information about the precise cellular source of SPARC and the developmental stage of cardiac progenitor cells responding to this signal.

**Fig. 1.**
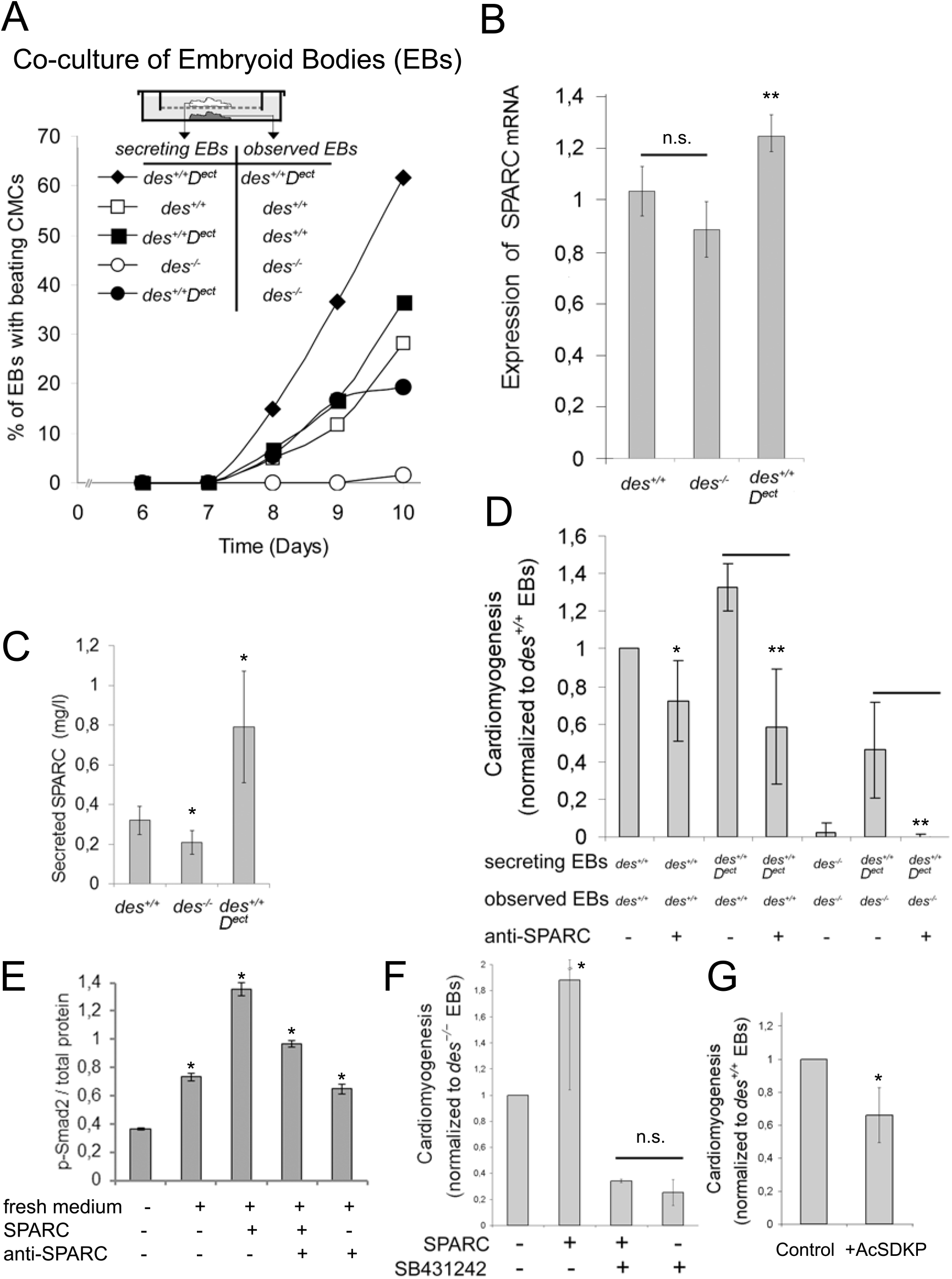
Increased desmin expression in EBs rescues early cardiomyogenesis in adjacent *des^-/-^* EBs in a paracrine manner via increased secretion of SPARC. **A**, Co-culture of desmin-overexpressing EBs (*des^+/+^D^ect^*) with wild-type (*des^+/+^*), or desmin-null (*des^-/-^*) EBs on permeable tissue culture sieve inserts promoted cardiomyogenesis in wild-type (*des^+/+^*) and *desmin* null (*des^-/-^*) EBs. The onset of cardiomyogenesis in EBs was monitored from day 6 to day 10 and the percentage of EBs with rhythmically contracting cardiomyocytes was determined. **B**, Desmin promotes SPARC expression in ESCs. Quantification of SPARC mRNA in ESCs of different genotypes by RT-sqPCR analysis. Number of experiments, n=3. GAPDH was used as reference gene. Error bars, standard deviation. **C**, Desmin promotes the secretion of SPARC from EBs. The concentration of secreted SPARC was calculated from dot blot analysis of media, used to culture EBs, with anti-SPARC antibodies and known concentrations of recombinant SPARC n=3. **D**; Sequestration of SPARC in the media with neutralizing antibodies attenuates the positive paracrine effect of *des^+/+^D^ect^*EBs on cardiomyogenesis in *des^+/+^* and *des^-/-^*EBs, respectively. EBs of the indicated genotypes (observed EBs) were cultured under sieve inserts with EBs used to secrete desmin-induced paracrine factors promoting cardiomyogenesis (secreting EBs), anti-SPARC antibodies were added from day 4.8 to day 7 of the in vitro differentiation experiment, and cardiomyogenesis was quantified between days 7 and 13 and normalized to the values obtained from *des^+/+^* EBs. **E**, Paracrine SPARC increases Smad2 phosphorylation in *des^-/-^* EBs, within 2 hours. Quantification of pSmad2 relative to total protein by Western blots. n=2. **F**, Promotion of cardiomyogenesis in *des^-/-^* EBs by SPARC is mediated by the TGF-ß receptor ALK5. ALK5 activity was inhibited by 3.5μmol/l SB431542in the presence or absence of 3mg/l SPARC for 48 hours. Cardiomyogenesis was normalized to that of *des^-/-^* EBs. **G**, Attenuation of nuclear import of pSmad2 by AcSDKP in *des^+/+^* EBs for 3 days partially suppressed cardiomyogenesis. (**A**, **D**, **F** and **G**), n=2; N (EBs)=20 per measurement; error bars, standard deviation; *, Student′s t-test p values p<0.05; **, p<0.01; n.s., not significant.

### Cardiac stem cells express and secrete SPARC and take up paracrine SPARC upon differentiation

SPARC is also expressed in the adult heart ^45^ (NCBI Gene ID: 6678) and protects the heart after injury ^46^, whereas knockout of *sparc* in mice negatively affects cardiac homeostasis ^8, 32^. To test whether SPARC may contribute to the role of adult CSCs and whether CSCs may be a significant source of SPARC in the adult heart, we screened CSC lines isolated from newborn and adult mice ^12^, by using immunofluorescence microscopy and RT-qPCR for the presence of SPARC mRNA and protein, respectively. CSCs expressed SPARC (**Figure 2A**) which was evenly distributed on the surface of undifferentiated CSCs, and the level of SPARC mRNA (**Figure 2B**) and SPARC protein (**Figure 2C**) in self-renewing CSCs was significantly higher than in ESCs. In the heart of adult mice, the cells with the highest SPARC protein levels sparsely populated the edge of cardiac muscle fibers (**Figure 2D**, arrowheads). To test whether isolated CSCs could be a source of secreted SPARC in the heart, we generated CSC lines expressing a SPARC-mCherry fusion protein (*des^+/+^sparc-mCherry^+^*). It was localized in the cytoplasm and occasionally in the nuclei of self-renewing CSCs (**Figure 2E** and **2F**). In differentiating CSC-derived cells SPARC was found in the perinuclear area where the Golgi apparatus is located and in many vesicles, similar to what has been observed in other cell types ^47^ (**Figure 2G**). CSCs and CSC-derived CBs secreted SPARC-mCherry into the medium (**Figure 2H**), which also bound to the ECM produced by CBs (**Figure 2I**). Culturing wild-type CSCs in SPARC-mCherry conditioned medium showed that differentiating CSC-derived cells take up SPARC after a lag phase of 24h, which has previously been demonstrated in CHO cells ^48^, whereas self-renewing CSCs barely take up SPARC from the medium (**Figure 2J**). From these data we may conclude that CSCs have the capability to produce, secrete and re-uptake significant amounts of SPARC in vitro after initiation of myocardial differentiation.

**Fig. 2.**
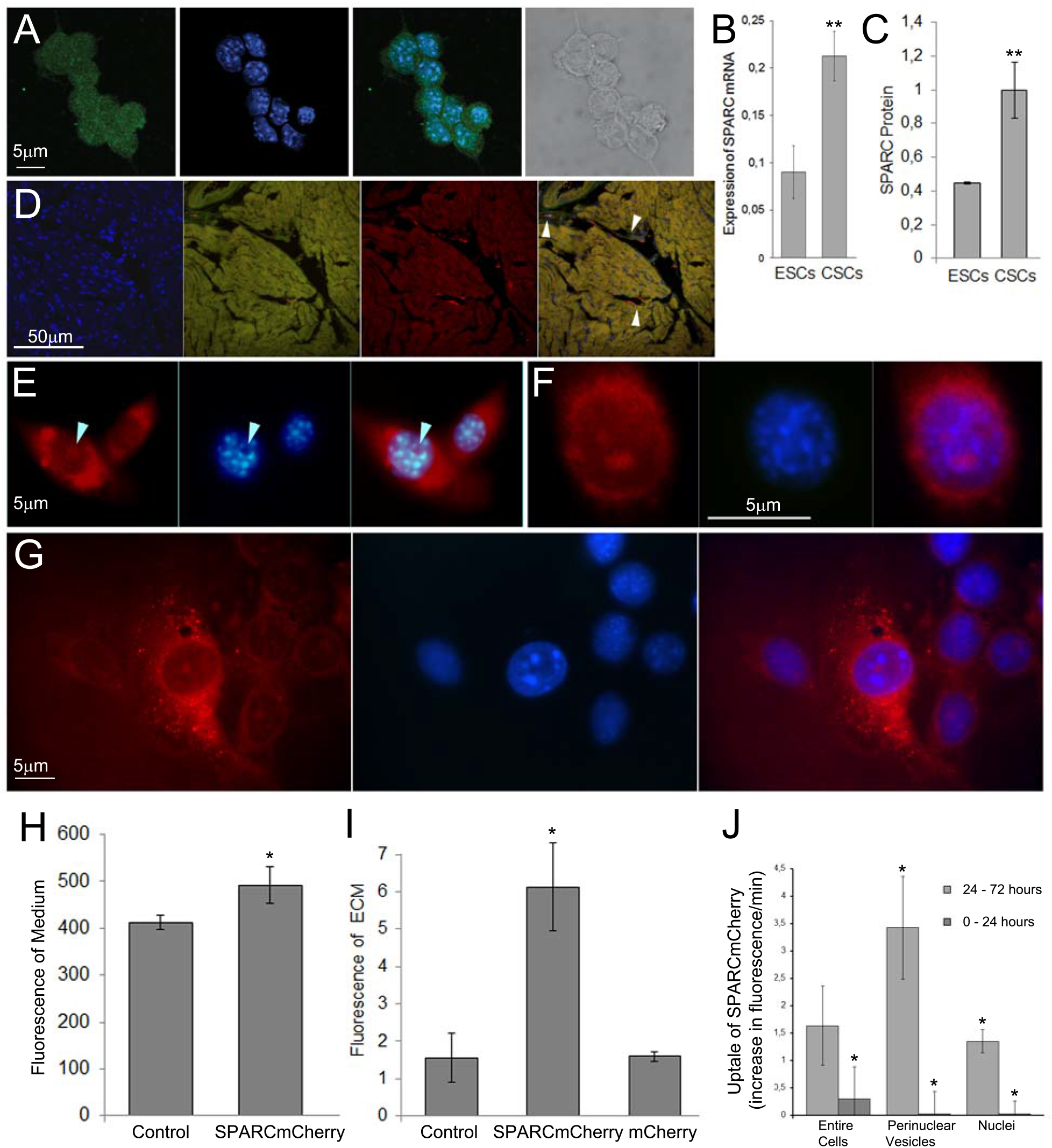
Cardiac stem cells express and secrete high levels of SPARC and incorporate SPARC as they differentiate to cardiac cells. **A**, Localization of SPARC on the surface of CSCs. Confocal immunofluorescence images of CSCs incubated with anti-SPARC antibodies (green) and DAPI-stained DNA (blue). **B**, RT-qPCR analysis of SPARC mRNA expression in CSCs compared to ESCs relative to RPL32 expression when cultured for 24 hours without feeder cells. n=5; error bars, standard deviation; **, Student′s t-test p value p=0.01. **C**, SPARC protein levels in CSCs and ESCs determined by quantitative Western blotting. N=2; p<0.001. **D**, Immunofluorescence images of mouse heart sections incubated with anti-SPARC antibodies (red), DAPI (blue) and anti-Sca1 (green) antibodies; Arrowheads in merged images, SPARC-positive cells at the edge of muscle fibers. **E**, Expression of SPARC-mCherry fusion protein in self-renewing CSCs. Fluorescence images showing expression of SPARC-mCherry (red), DAPI-stained DNA (blue). Representative images from 2 out of 10 transgenic cell lines showing cytoplasmic and nuclear localization of SPARC-mCherry protein (arrowheads). **F**, Perinuclear and nuclear localization of SPARC-mCherry in self-renewing CSCs. **G**, Cytoplasmic, vesicular and nuclear localization of SPARC-mCherry in differentiated CSC progeny cells. **H**, SPARC-mCherry fluorescence in media conditioned by SPARC-mCherry expressing CBs and wild-type CBs (control). Data are from 10 measurements. n=3; error bars, standard deviation; p=0.0001. **I**, SPARC-mCherry secreted by CBs but not mCherry, also secreted by CBs, binds to an ECM produced by wild-type CBs. Control, medium conditioned with wild-type CBs. Data are from n=3 independent experiments (SPARC-mCherry and control) and n=2 experiments (mCherry); error bars, standard deviation; p<10^-8^. Note, control levels in **H** and **I** represent autofluorescence of the medium and CSC derived ECM, respectively. **J**, Rate of SPARC-mCherry fusion protein uptake into differentiating CSCs, perinuclear vesicles and nuclei, respectively, between 0 and 24 hours and 24 and 72 hours, respectively, after initiation of differentiation by withdrawal of LIF. Data are from 4 measurements each. n=2; error bars, standard deviation; *, Student′s t-test p value p<0.05.

### Paracrine and autocrine SPARC promotes cardiomyogenesis in differentiating cardiac stem cells

To test whether *sparc* is expressed during in vitro cardiomyogenesis in CSC-derived cells, we induced differentiation by aggregating them in hanging droplets in the absence of LIF and feeder cells to form CBs ^12^ and quantified and compared SPARC mRNA expression by RT-qPCR over time with SPARC expression in ESC-derived EBs (**Figure 3A**). SPARC expression continuously increased with progression of differentiation in both cell types and it was constantly higher in CBs than in EBs, whereas the decrease in *nanog* expression was similar in CBs and EBs. The latter suggests that the progression of differentiation per se was comparable in CBs and EBs (**Figure 3B**). To test whether autocrine SPARC is necessary for cardiomyogenesis in CSCs and to determine the developmental stage that mainly depends on the presence of extracellular SPARC, we cultured CBs in the presence of SPARC-neutralizing antibodies for different time periods and compared cardiomyogenesis in these CBs with that of untreated CBs by quantifying the number of rhythmically contracting cardiomyocytes between day 13 and day 20 of CB development (**Figure 3C**). Reduced extracellular SPARC levels attenuated cardiomyogenesis in CBs during the first seven days of differentiation and did not significantly affect cardiomyogenesis later, between days 7 and 13. Addition of paracrine SPARC to CBs from day 4.8 to day 7, determined to be the most SPARC-sensitive developmental stage of cardiomyogenesis in vitro, accelerated cardiomyogenesis (**Figure 3D**) and increased the number of rhythmically contracting cardiomyocyte clusters by 32% (**Figure 3E**), whereas addition of SPARC-neutralizing antibodies delayed cardiomyogenesis (**Figure 3D**) and reduced the number of rhythmically contracting cardiomyocyte clusters to 42% of the control (**Figure 3E**). Next we asked whether SPARC produced by CSCs influences cardiomyogenesis in CBs in a dose-dependent manner. Therefore, we generated CBs from heterozygous *sparc* knockout CSC lines (*sparc^+/-^*), as well as from CSC lines additionally expressing *sparc* under the control of a CMV promoter from an ectopically integrated *sparc^ect^* transgene (*sparc^+/+^S^ect^*) and compared cardiomyogenesis in these CBs with cardiomyogenesis in *sparc^+/+^*wild-type A5 (**Figure 3F**). Increased expression of *sparc* had a pronounced positive effect on the initial increase and extent of cardiomyogenesis. In contrast, deletion of one *sparc* allele resulted in a haploinsufficiency phenotype manifested by a delay in cardiomyogenesis and a reduced number of cardiomyocytes in *sparc^+/-^* CSC-derived CBs. Similarly, expression of SPARC-mCherry from an ectopically integrated *sparc::mCherry^ect^*transgene (*sparc^+/+^S:mC^ect^*), which is also secreted into the medium (see **Figure 2F**), significantly accelerated cardiomyogenesis in CBs compared to mCherry-expressing (*sparc^+/+^mC^ect^*) CBs (**Figure 3G**). Reducing the density of CBs on the plates and thus the concentration of autocrine secreted wild-type SPARC in the medium demonstrated the acceleration of cardiomyogenesis by SPARC-mCherry even more clearly (**Figure 3H**). The frequency of rhythmic contraction of *sparc^+/+^S^ect^* cardiomyocytes was significantly increased when compared to wild-type cardiomyocytes and diminished when anti-SPARC antibodies were added to the medium (Supplemental Figure **1A**). These data are consistent with the observation that SPARC induces a positive inotropic effect in murine hearts ^49^.

**Fig. 3.**
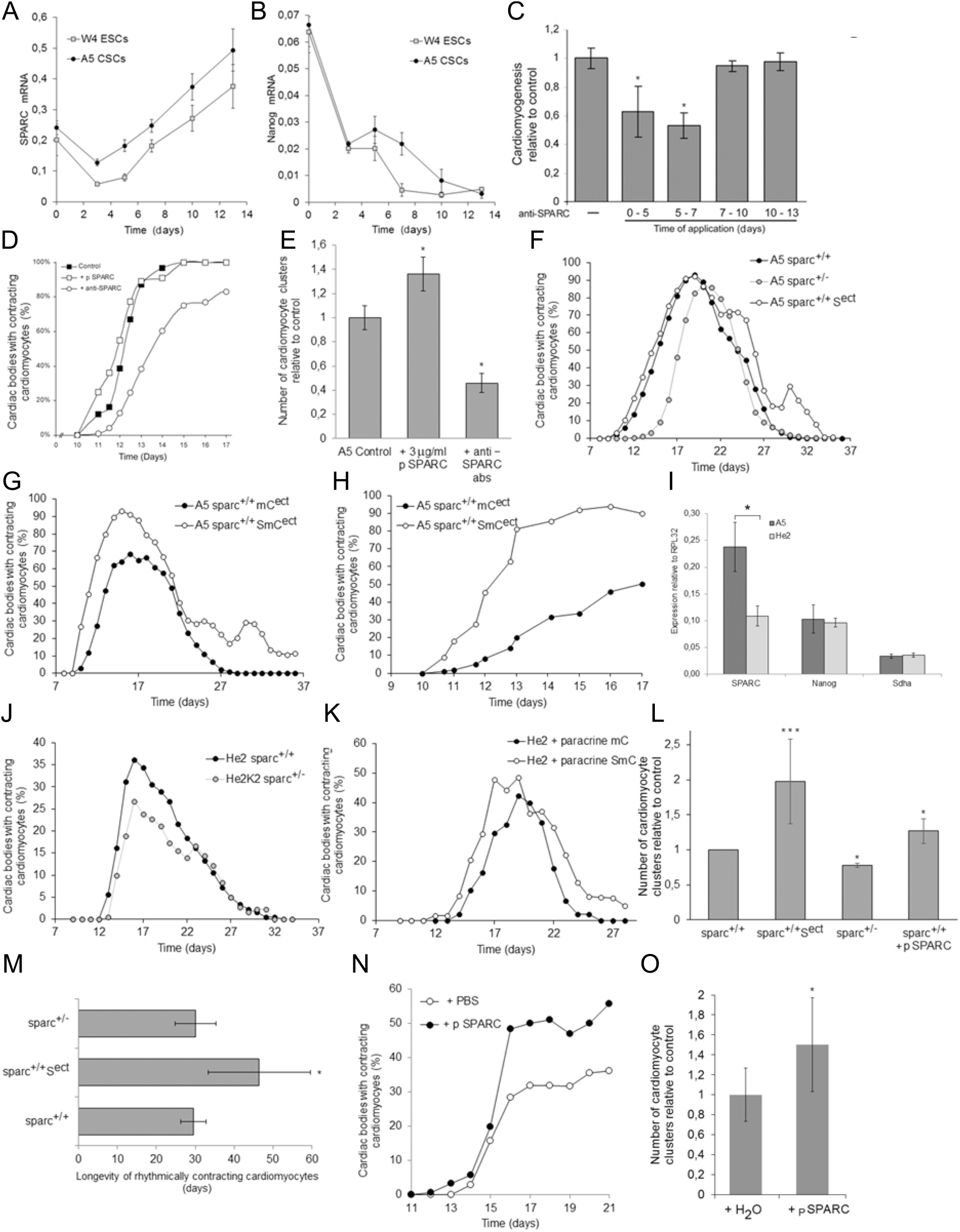
SPARC positively influences cardiomyogenesis in a paracrine and autocrine dose-and developmental-stage dependent manner. **A**, Expression of SPARC and **B**, nanog mRNA in developing A5 wild-type CSC-derived CBs and W4 wild-type ESC-derived EBs after withdrawal of feeder cells and aggregation at d0. RT-qPCR analysis using primer pairs for SPARC and nanog and RPL32, as internal reference. Data from 3 biological replicates and 2 technical replicates each; number of experiments n=3. Error bars, standard deviation; Student′s t-test p values for all SPARC and the nanog day 7 data, p<0.05. **C**, Inhibition of extracellular autocrine SPARC negatively affects cardiomyogenesis in A5 wild-type CBs between days 0 and 7 before cardiomyocytes start to contract at day 11. Anti-SPARC antibodies were added to CBs for time intervals as indicated on the x-axis. Cardiomyogenesis was assessed by counting the number of beating clusters of cardiomyocytes per CB between days 13 and 20. Data are normalized to control without anti-SPARC antibodies. Error bars, standard deviation; Student′s t-test p values for day 0-5 and 5-7 data, p<0.05. **D**, Surplus paracrine SPARC accelerates cardiomyogenesis between days 10 and 14 in A5 CBs, whereas neutralizing antibodies against SPARC significantly delay cardiomyogenesis. Time course of myocardial differentiation in CBs. Data represent the percentage of CBs with at least one rhythmically contracting cluster of cardiomyocytes. Paracrine recombinant SPARC (2mg/l) (pSPARC) and neutralizing amounts of anti-SPARC antibodies, respectively, were added between days 4.8 and 7. **E**, Paracrine SPARC increases the number of cardiomyocyte clusters per CB, and inhibition of SPARC by neutralizing antibodies reduces the number of cardiomyocyte clusters. SPARC was added to CBs between days 4.8 and 7 and the number of cardiomyocyte clusters in CBs was determined between days 9 and 13. (**D** and **E**) n=2; 3 biological replicates each. Error bars, standard deviation; *, p<0.05. **F**, Overexpression of SPARC increases the onset of cardiomyogenesis and the percentage of CBs with contracting cardiomyocytes, whereas mono-allelic expression delayed and attenuated cardiomyogenesis in CBs. Development of contracting cardiomyocytes in CBs over time. *sparc^+/+^,* wild-type CSCs; *sparc^+/+^S^ect^*, SPARC-overexpressing CSCs (mean of 2 cell lines A5K4 and A5K10); *sparc^+/-^,* heterozygous knockout cell lines (mean of 2 cell lines A5K11 and A5K17). Data from each cell line from 3 technical replicates; n=2; N (CBs)=60 each. **G**, Overexpression of SPARC-mCherry fusion protein increases the onset of cardiomyogenesis, the number of contracting cardiomyocytes, and the longevity of contracting cardiomyocytes compared to mCherry-expressing CBs. Development of contracting cardiomyocytes in CBs over time. *sparc^+/+^S:mC^ect^*, SPARC-mCherry-expressing CBs; *sparc^+/+^mC^ect^*mCherry-expressing CSCs. Data from 3 technical replicates; n=2; N (CBs)=60 each. **H**, Reducing the time intervals for monitoring the onset of myocardial contraction in CBs between day 10 and day 13 and the density of CBs on a plate demonstrates a significant acceleration of cardiomyogenesis by SPARC-mCherry fusion protein. Other experimental conditions as in 3G. **I**, Expression of SPARC mRNA in A5 CSCs and He2 CSCs. RT-qPCR analysis with primer pairs for SPARC, nanog, and sdha, the latter two as internal controls. Data from 3 biological replicates with each 2 technical replicates. Error bars, standard deviation; *, p<0.05. **J**, Mono-allelic expression of SPARC (*sparc^+/-^*) in He2K2 CSC-derived CBs with a Balb/c background delays and reduces cardiomyogenesis more severely than in A5 wild-type CBs. Experimental conditions as in 3G. Note that the onset of rhythmic contraction of cardiomyocytes in He2 CBs with a Balb/c background is delayed by 1 day and by 2 days in He2K2 compared to A5 CBs. **K**, Paracrine addition of secreted SPARC-mCherry fusion protein to wild-type He2 CBs in co-cultures with A5 *sparc^+/+^S:mC^ect^* CBs (He2 + paracrine SmC) also increases the rate of cardiomyogenesis, affects the onset of cardiomyogenesis, and increases the longevity of rhythmically contracting cardiomyocytes compared to He2 CBs co-cultured with A5 *sparc^+/+^mC^ect^*control CBs (He2 + paracrine mC). Experimental conditions as in 3G. **L**, Mean number of contracting cardiomyocyte clusters per CB monitored from day 13 to day 26. Data are from experiments shown in **3C**–**3H**. Error bars, standard deviation; ***, p<0.001; *, p<0.05. **M**, Longevity of rhythmically contracting cardiomyocytes in CBs ectopically expressing SPARC and with mono-allelic expression of SPARC compared to wild-type cardiac bodies. Data are from n=3 or more experiments (**3C**-**3H**) and n=2 (3J and 3K) independent experiments with 3 technical replicates each. **N**, Addition of 3mg/l paracrine recombinant SPARC from day 4.8 to 7 rescues cardiomyogenesis in *sparc^+/-^* CBs and, **O**, increases the number of clusters with contracting cardiomyocytes. Data are from 3 technical replicates; n=2; N (CBs)=60 each. Mean number of individual contracting cardiomyocyte clusters per CB monitored from day 16 to day 19. Error bars, standard deviation; *, Student′s t-test p value p<0.05.

To further extend the range of autocrine SPARC expression, we performed similar experiments with He2 CSC lines with a *Balb/c* genetic background which express significantly less SPARC than the A5 CSC lines derived from a *C57BL/J6:129Sv* genetic background ^12^ (**Figure 3I**). Indeed, the negative effects of the loss of one *sparc* allele observed in the *C57BL/J6:129Sv* genetic background was even more pronounced in the *Balb/c* genetic background in *sparc^+/-^* He2K2 CBs (**Figure 3J**). To test whether paracrine SPARC-mCherry can ameliorate cardiomyogenesis in He2 CBs and to see whether it is indeed the low SPARC level in He2 CBs that causes the lower cardiomyogenic potential, we co-cultured He2 CBs in the presence of SPARC-mCherry A5 CBs or mCherry expressing A5 CBs, respectively, as control (**Figure 3K**). The presence of SPARC-mCherry conditioned medium again accelerated the progression of myocardial differentiation, and increased the extent and duration of cardiomyogenesis, and the longevity of cardiomyocytes in He2 CBs. Comparing the mean number of rhythmically contracting cardiomyocyte clusters in A5 CBs with different numbers of *sparc* alleles between days 13 and 26 demonstrates that autocrine overexpression of *sparc* nearly doubled cardiomyogenesis (190%), mono-allelic expression of *sparc* reduced cardiomyogenesis to 80% of control, and paracrine addition of recombinant SPARC increases cardiomyogenesis by 30% compared to control (**Figure 3L**). Long-term experiments showed that autocrine overexpression of *sparc* in A5 and He2 CBs significantly increased longevity and robustness of rhythmically contracting cardiomyocytes (**Figure 3M**). Mono-allelic expression of *sparc* delayed and reduced cardiomyogenesis in both genetic backgrounds (Figure **3F** and **3J**), but did not reduce the longevity of rhythmically contracting cardiomyocytes (**Figure 3M**). Culturing primary mouse cardiomyocytes on CSC-derived feeder cells ectopically expressing more *sparc* prolonged the survival and the rhythmic contraction of these cells (Supplemental Figure **1B** and **1C**) and doubled the contraction frequency (Supplemental Figure **1D**), as has been previously reported ^49^. Finally, addition of 3mg/l paracrine SPARC from day 4.8 to day 7 partially rescued the haploinsufficiency in *sparc^+/-^* CBs, both by promoting cardiomyogenesis (**Figure 3N**) and by increasing the number of clusters with rhythmically contracting cardiomyocytes (**Figure 3O**). Taken together, these results suggest, first, that CSCs and their differentiating progeny may be an important source of SPARC in the heart, second, that SPARC strongly promotes cardiomyogenesis but is most likely neither necessary nor sufficient to induce cardiomyogenesis in vitro, and third, that both autocrine and paracrine SPARC contribute to cardiomyogenic differentiation of CSCs in a dose-and developmental stage-dependent manner.

### Paracrine SPARC influences gene expression already in self-renewing cardiac stem cells

The promotion of cardiomyogenesis by paracrine and autocrine SPARC raises the question whether its beneficial effect is mediated by its influence on the expression of mesodermal or myocardial transcription factor genes in CSCs. In subconfluent cultures of CSCs addition of 3mg/l recombinant SPARC increased mRNA expression of primitive mesodermal transcription factor genes *brachyury* (*T*) and *goosecoid* but did not influence the expression of *isl1* and *mef2C* (**Figure 4A**). The early myocardial transcription factor genes *mesp1*, *nkx2.5* and g*ata4*, as well as the very early myocardial marker *desmin* ^10^ were upregulated within 2 hours, whereas *cTnT* expression, as expected, was not affected (**Figure 4B**).

**Fig. 4.**
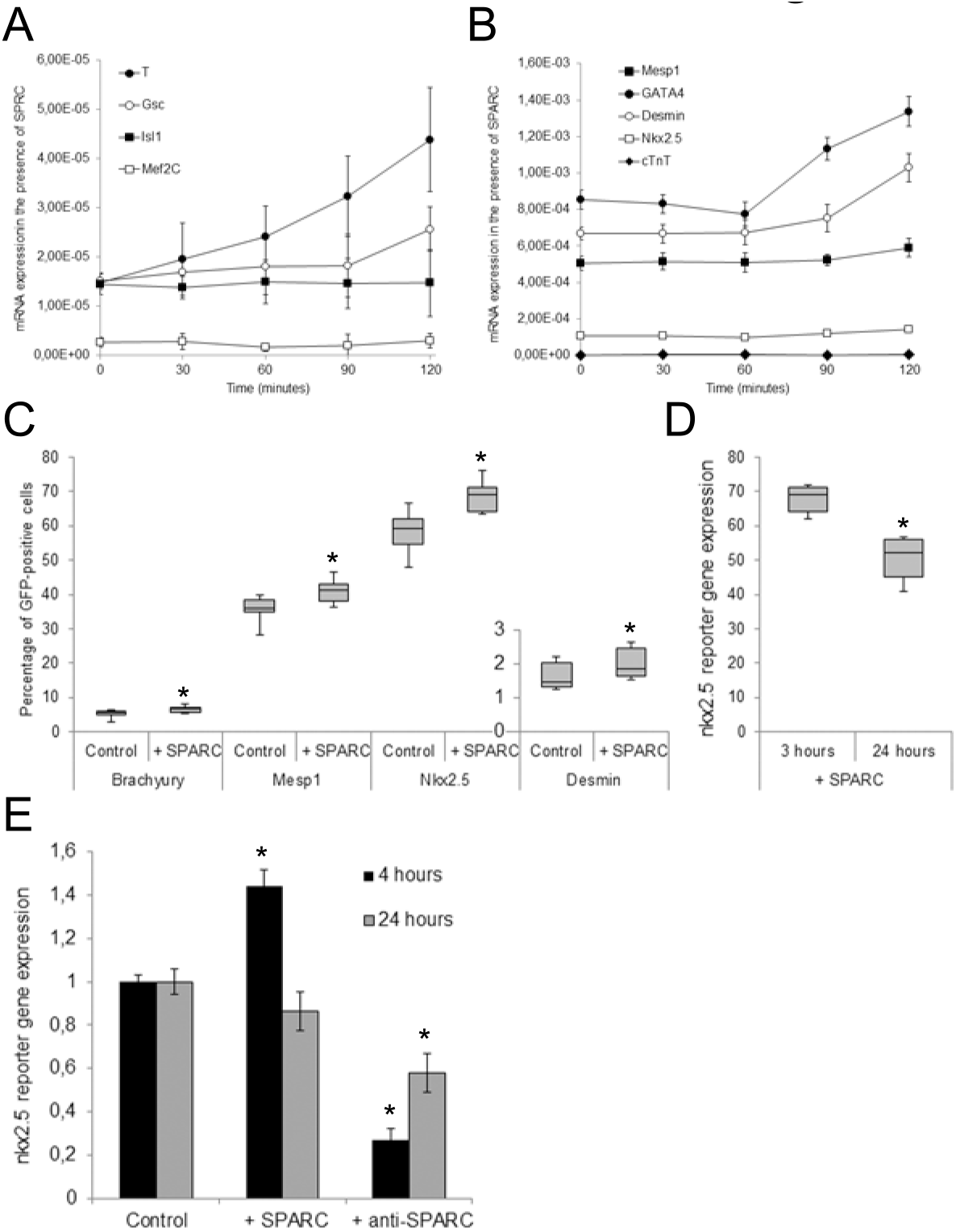
Paracrine SPARC promotes expression of early mesodermal and myocardial transcription factors and of desmin. **A** and **B**, RT-qPCR analysis of mRNA expression of indicated genes in CSCs in the absence of LIF and 3mg/l recombinant SPARC. n=2, 2 biological and 2 technical replicates each. mRNA levels were normalized to RPL32 mRNA. **C**, Fluorescence activated cell sorting analysis of the percentage of eGFP-positive CSCs expressing a *brachyury*, *mesp1*, *nkx2.5*, or a *desmin* eGFP reporter gene in the presence of conditioned medium from STO and 3T3 fibroblasts (control) or STO and 3T3 fibroblasts secreting SPARC from an expression cassette, after 3 hours of incubation. Box plots show the median and the second and third quartiles; whiskers indicate maximum and minimum values, respectively. n=3; **D**, Expression of an *nxk2.5::eGFP* knock-in allele in CSC after incubation with medium from SPARC-secreting STO cells for 3 and 24 hours. Whiskers blot as in B. **E**, Expression of an *nkx2.5::luciferase-reporter* gene in transiently transfected, CSC-derived, differentiated cardiac cells from 9-day-old CBs. Fluorescence measurements were performed 24 hours after transfection plus 4 or 24 additional hours of incubation with medium from SPARC-secreting STO cells. n=3; error bars, standard deviation; *, all Student′s t-test p values p<0.05.

To obtain additional evidence at the single cell level that SPARC contributes to the myocardial differentiation of CSCs by influencing transcription, we introduced eGFP-expressing reporter genes for *T* ^40^, *mesp1* ^41^ and a *desmin::EGFP* reporter gene containing 3.4 kb of the *desmin* enhancer and promoter region by non-homologous recombination. A knock-in *nkx2.5* reporter construct ^42^ was introduced by homologous recombination. We isolated clonal cell lines expressing these reporter genes and measured the percentage of CSCs with increased eGFP fluorescence by fluorescence activated cell sorting in the presence of SPARC-secreting STO or 3T3 fibroblasts or wild-type STO or 3T3 fibroblasts, which were used as feeder cells to maintain CSC self-renewal (**Figure 4C**). SPARC indeed induced expression of *T*, *mesp1*, *nkx2.5*, and *desmin* in a subset of self-renewing CSCs. The fact that the percentage of cells expressing these reporter genes varied greatly suggests that expression, even in the absence of SPARC, was stochastic in nature and would be channeled to a particular developmental cue only later during myocardial differentiation of CSCs. However, presence of paracrine SPARC significantly increased the number of cells expressing the reporter genes and thus by these means may contribute to the promotion of cardiomyogenesis during CB development, as shown in Figure **3D** and **3E**. In addition to this apparently incomplete penetrance of SPARC signaling in self-renewing CSCs, paracrine SPARC also resulted in only a short-term activation of the *nkx2.5* gene, as previously demonstrated for *T* ^50^ and *mesp1* ^51^ during mesoderm development in EBs. Maintenance of CSCs in SPARC-containing medium and on feeder cells for 24 hours resulted in a reduced number of CSCs expressing the *nkx2.5::eGFP* reporter gene (**Figure 4D**). Short-term but not long-term induction of *nkx2.5* expression by SPARC was also confirmed by measuring the luciferase activity in differentiating CSCs transiently transfected with a *nkx2.5::Luciferase* reporter construct containing only the proximal promoter region of the *nkx2.5* gene ^44^ (**Figure 4E**). Conversely, the activation of the reporter genes was inhibited by anti-SPARC antibodies.

The induction of *T*, *mesp1*, *nkx2.5*, and *desmin* expression in only a subset of CSCs and its transient nature suggests that either the available reporter genes do not contain all regulatory elements necessary for in vivo-like gene expression in CSCs or that SPARC protein levels above a certain threshold may be necessary but not sufficient for proper myocardial gene expression during the developmental stages at the onset of cardiomyogenesis. Nevertheless, SPARC seems to influence myocardial differentiation in CSCs from the very beginning, by modulating the expression of early myocardial transcription factors and of *desmin*. The latter, together with the fact that desmin promotes *sparc* expression in differentiating ESCs, see **Figure 1B**, raises the question of whether changes in the basal protein levels of SPARC and desmin in CSCs also influence the mRNA expression of desmin and SPARC and thus the developmental fate of CSCs.

### SPARC and desmin alternately promote their mRNA expression and protein levels in CSCs at the onset of differentiation

Desmin was expressed at significantly higher levels in CSCs than in ESCs, and concomitantly, CSCs expressed and secreted more SPARC than ESCs, see Figure **2B** and **3C**. In addition, moderately increased expression of *desmin* in ESCs led to an increased expression and secretion of SPARC and to increased cardiomyogenesis in EBs ^15, 18^, however, in this model system it was not possible to distinguish a possible cell-autonomous effect in cardiac progenitor cells from the obvious non-cell-autonomous effects mediated by paracrine SPARC also secreted by extra-embryonic endoderm ^14, 52^, and the obvious mutual interaction between different types and populations of cells in EBs. To test the hypothesis that not only SPARC promotes *desmin* expression, but that desmin also promotes SPARC expression and secretion in CSCs, and further that desmin contributes to decision making during CSC specification, we generated *des^+/-^* and *des^+/+^D^ect^* CSC lines and used the previously generated *des^+/+^D:mC^ect^* cell lines ^16^ and compared SPARC and desmin mRNA expression and protein levels in them with those in *sparc^+/-^* and *sparc^+/+^S^ect^* CSC lines, respectively. As expected, SPARC mRNA expression and protein synthesis were altered according to the genotype of mutant *sparc* CSC lines (Figure **5A** and **5B**) and desmin mRNA and protein levels correlated with the SPARC mRNA and protein levels (Figure **5C** and **5D**). In turn, mutant *desmin* CSC lines expressed less or more desmin mRNA and protein than the wild-type CSCs (Figure **5E** and **5F**), and again SPARC mRNA and protein levels correlated with the desmin mRNA and protein levels of mutant *desmin* CSC lines (Figure **5G** and **5H**). These data suggest a cell-autonomous correlative influence of the dosage of SPARC expression on *desmin* expression and vice versa and support the hypothesized positive genetic feedback loop between *sparc* and *desmin* in CSCs. Furthermore, mono-allelic expression of *sparc* and *desmin*, respectively, resulted in a reduced secretion of SPARC, and *sparc*-and *desmin*-overexpressing CSC lines secreted more SPARC into the cell culture supernatant than wild-type CSCs (**Figure 5I**). Finally, addition of paracrine SPARC to wild-type, *des^+/-^*, and *sparc^+/-^* CSCs and CBs, respectively, resulted in a downregulation of SPARC mRNA and protein expression (**Figure 5J and K**), which adds the possibility that the positive SPARC-desmin loop can be muted by a paracrine negative SPARC-*sparc* feedback loop. These data together suggest that differential expression of *desmin* also contributes to the autocrine regulation of CSC differentiation by affecting the levels of translated and secreted SPARC protein.

**Figure 5.**
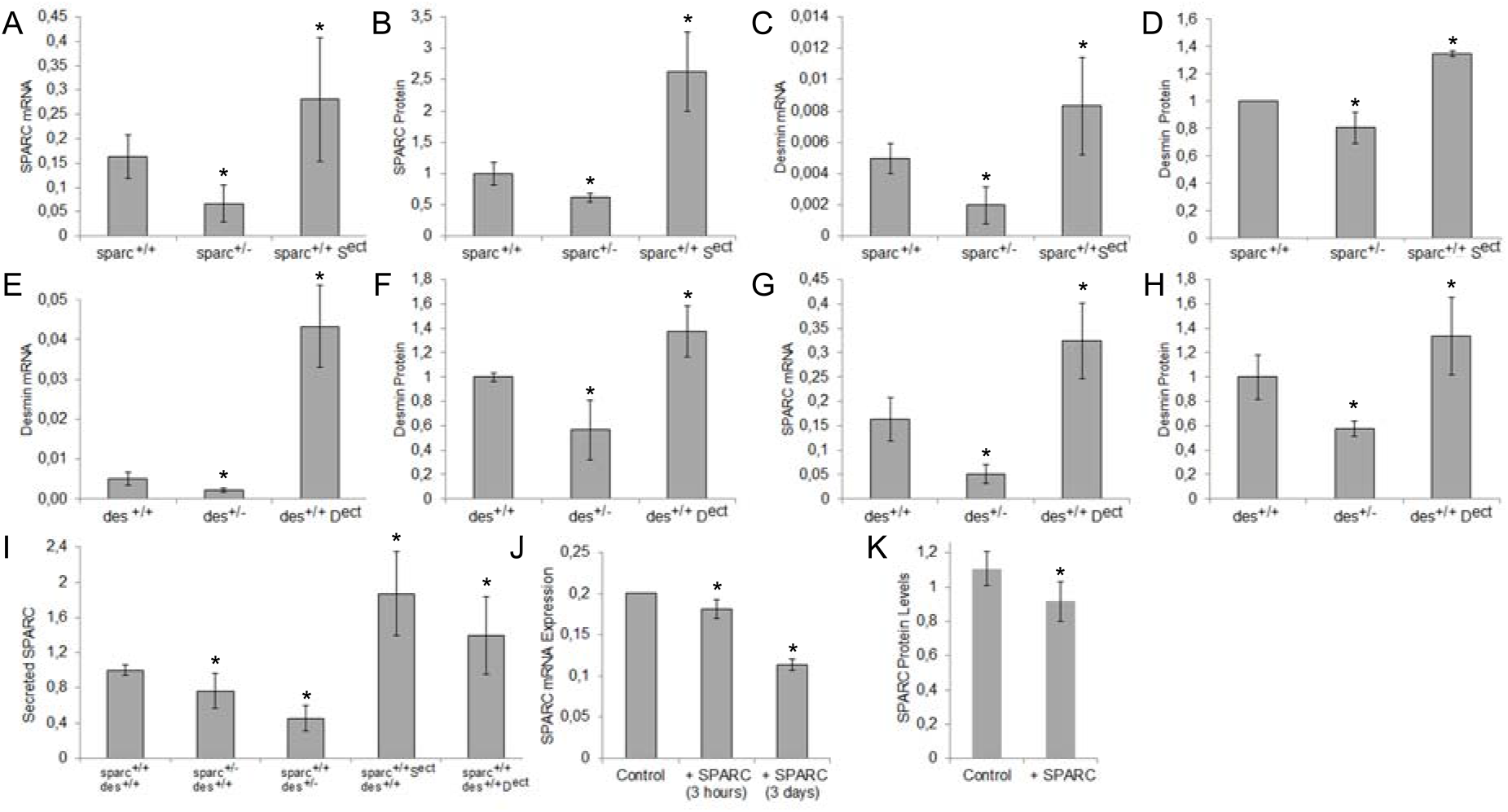
SPARC and desmin alternately promote each otheŕs mRNA expression and protein levels in CSCs at the onset of differentiation. SPARC and desmin mRNA expression levels and cytoplasmic protein concentrations were quantified in CSC lines with mono-allelic expression of either desmin or SPARC, and in CSC lines ectopically expressing desmin or SPARC in addition to the bi-allelic expression. mRNA and protein quantities were compared to the parental wild-type *sparc^+/+^ des^+/+^* CSC line. mRNA levels were normalized to RPL32 mRNA levels and protein levels were normalized to total protein found in the same number of cells of each CSC line. **A**, SPARC mRNA levels and **B**, SPARC protein levels in 2 *sparc^+/-^* and 2 *sparc^+/+^S^ect^*CSC lines. **C**, Desmin mRNA levels and **D**, desmin protein levels in the same *sparc^+/-^* and *sparc^+/+^S^ect^* CSC lines. **E**, Desmin mRNA levels and **F**, Desmin protein levels in each 3 *des^+/-^* and 2 *des^+/+^D^ect^* CSC lines, respectively. **G**, SPARC mRNA levels and **H**, SPARC protein levels in the same *des^+/-^*and *des^+/+^D^ect^* CSC lines. Number of independent experiments, n≥3; all Student′s t-test p values p≤0.001. **I**, Quantification of SPARC protein secreted into the cell culture supernatant of *sparc* and *desmin* mutant CSC lines, respectively. Protein concentration in the culture medium was normalized to that of total protein found in CSC cultures with the same number of cells after 24 hours of culture. n≥3; all Student′s t-test p values p<0.05. **J**, SPARC mRNA expression in wild-type, *sparc^+/-^*and *des^+/-^* CSCs 3 hours after addition of 3mg/l SPARC and in CBs 3 days after SPARC addition. **K**, SPARC protein levels (arbitrary units) in CSCs after addition of SPARC for 3 hours; conditions as in J. Number of independent experiments, n=2; with 2 technical replicates each; all Student′s t-test p values p≤0.003.

### Reduced and increased expression of desmin in CSCs negatively affects cardiomyogenesis

To test whether altered expression and secretion of SPARC contributes to the cardiomyogenic phenotype of *des^+/-^* and *des^+/+^D^ect^*CBs, we first monitored cardiomyogenesis in CBs generated from these cell lines. Consistent with decreased desmin and SPARC protein levels in these cells, cardiomyogenesis was reduced and attenuated in *des^+/-^* CBs (Figure **6A** and **6B**), similar to what was observed in *sparc^+/-^* CBs (Figure **3F** and **3J**) and *des^+/-^*EBs ^37^. However, in contrast to *desmin*-overexpressing EBs ^15^ and *sparc*-overexpressing CBs (**Figure 3F-H** and **3K**), desmin and desmin-mCherry overexpression above what could be considered physiologically meaningful levels in *des^+/+^D^ect^* CSCs resulted in reduced cardiomyocyte development and shortened the lifespan of cardiomyocytes, despite increased SPARC secretion (**Figure 5A**). Desmin overexpression also caused an increase in beating frequency and mono-allelic expression a decrease in beating frequency (**Figure 6C**), as already observed in EBs ^15^. However, here in CSC-derived cardiomyocytes this was accompanied by a severe arrhythmia at the cellular level (**Figure 6D**), probably causing attenuation of cardiomyogenesis and premature death of cardiomyocytes. Similarly, cardiomyogenesis was impaired in desmin-mCherry fusion protein expressing CSC-derived CBs (**Figure 6A** through **6D**). In these cells, massive overexpression of *desmin* led to its aggregation in some clusters of cardiomyocytes (**Figure 6E**), likely contributing to arrhythmia and the inability to contract for prolonged periods of time. In contrast, mCherry expression alone did not induce arrhythmic contractions of cardiomyocytes (see **Figure 3G**) and was homogeneously and diffusely distributed throughout the entire cell (not shown). Apart from the dominant negative effect of too much desmin in developing cardiac cells, which might obscure of the positive effect of SPARC seen in *sparc^+/+^ S^ect^* CBs, these data additionally support the hypothesis that moderate expression of *desmin* also contributes to the autocrine regulation of CSC differentiation by influencing the levels of translated and secreted SPARC.

**Figure 6.**
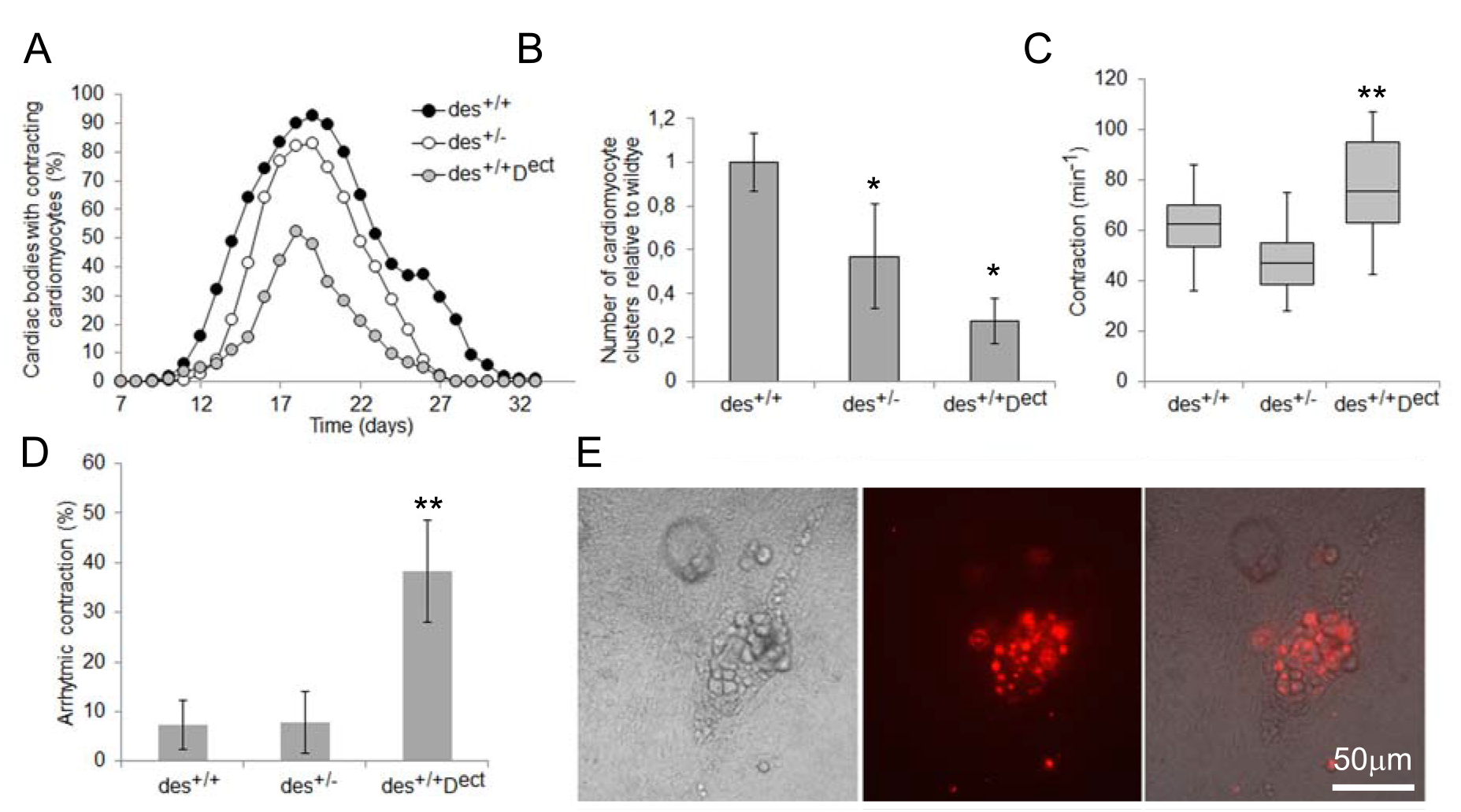
Reduced and increased expression of desmin negatively affects cardiomyogenesis in CSCs. Cardiomyogenesis in CBs with either mono-allelic or overexpression of desmin. **A**, Time course of cardiomyogenesis in wild-type *des^+/+^*, *des^+/-^*, and *des^+/+^D^ect^*, *des^+/+^D:mC^ect^*, and *des^+/+^mC:D^ect^* CBs. Data were collected from 4 *des^+/-^*and 3 *des^+/+^D^ect^* cell lines as well as from 1 *des^+/+^D:mC^ect^*and 1 *des^+/+^mC:D^ect^* cell line, each; number of experiments, n=3; N (CBs)=60 from 3 technical replicates each;. **B**, Mean number of contracting cardiomyocyte clusters per CB. Data were collected between days 14 and 18 of CB development; n=3; N=60, each. **C**, Frequency of rhythmic contraction of cardiomyocyte clusters in CBs. Box plots show the median and the second and third quartiles; whiskers indicate maximum and minimum values, respectively. Data were collected between days 11 and 26 of CB development. n=3; each N≥60; *, Student′s t-test p value p<0.05. **D**, Percentage of clusters with arrhythmically contracting cardiomyocytes. **, Student′s t-test p value p<0.01. **E**, Phase contrast and fluorescence microscopy images of an arrhythmically contracting cardiomyocyte cluster with desmin aggregates in *des^+/+^D:mC^ect^* CBs. Bar, 50 μm.

### Paracrine glycosylated SPARC rescues the cardiomyogenic haplo-insufficiency phenotypes in *des^+/-^* cardiac bodies and promotes expression of myocardial transcription factor genes

Correlation of the level of secreted SPARC with the amount of cardiogenesis in differentiating CSC lines and correlation with the expression levels of both the *sparc* and *desmin* genes suggest that secreted SPARC may partially rescue haploinsufficiency in *des^+/-^* CSC lines. To test whether SPARC must be post-translationally modified and secreted, or if intrinsic, cytoplasmic SPARC suffice to exert its effects on cardiomyogenesis and to rescue *desmin* haploinsufficiency, we added glycosylated recombinant SPARC and de-glycosylated SPARC to *des^+/-^*CBs between day 4.8 and 7.5 and monitored cardiomyogenesis over time. Paracrine administration of glycosylated SPARC (**Figure 7A**) rescued cardiomyogenesis compared to the administration of de-glycosylated SPARC (**Figure 7B**) and significantly increased the number of cardiomyocyte clusters in CBs (**Figure 7C**), whereas de-glycosylated SPARC had only a small, insignificant positive effect (**Figure 7D**), possibly due to the incomplete de-glycosylation or endocytic uptake. Glycosylated SPARC also had a positive effect on the EMT at the onset of cardiomyogenesis in CBs from day 4.8 to 6.7 (**Figure 7E**), whereas de-glycosylated SPARC did not (**Figure 7F**). The frequency of rhythmic contraction was increased by glycosylated SPARC but not by de-glycosylated SPARC (**Figure 7G**). Finally, addition of glycosylated SPARC to CSCs and CBs for 3 hours and 3 days, respectively, or overexpression of *sparc* in CSCs increased the pSmad2 to Smad2 ratio (**Figure 7H**). This suggests, as already shown for EBs (see Figure 1E), that SPARC influences Tgf-ß-Smad2 signaling also in CSCs and their differentiating progeny.

**Figure 7.**
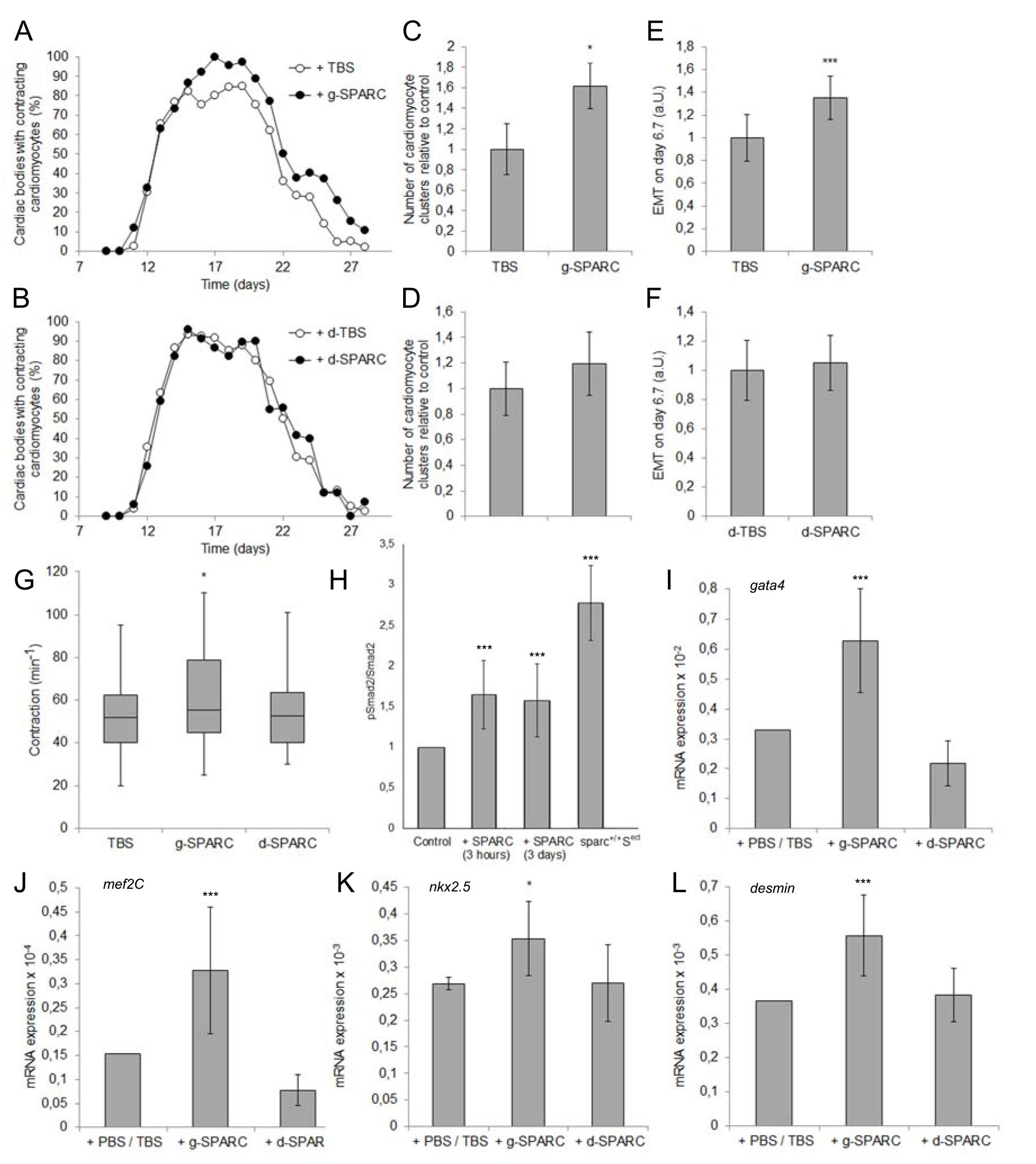
SPARC rescues cardiomyogenesis in *des^+/-^* CBs in a glycosylation-dependent manner and promotes the expression of cardiomyogenic transcription factor genes. Cardiomyogenesis in CBs with either mono-allelic expression of desmin or SPARC in the presence of either glycosylated SPARC (**A**, **C** and **E**) or de-glycosylated SPARC (**B**, **D** and **F**). **A** and **B**, Time course of cardiomyogenesis in *des^+/-^* CBs. Data were collected from 2 *des^+/-^*cell lines in 2 biological and 3 technical replicates; number of experiments n=3; N (CBs)=20 each. **C** and **D**, Mean number of contracting cardiomyocyte clusters per CB. Data were collected between days 14 and 18 of CB development; n=2; N=60 from 3 technical replicates each. **E** and **F**, EMT and migration of cells in CBs measured as the increase of the diameter of CBs in 24h. **G**, Frequency of rhythmic contraction of cardiomyocyte clusters in CBs. Box plots show the median and the second and third quartiles; whiskers indicate maximum and minimum values, respectively. Data were collected between days 12 and 22 of CB development. n=2; N≥60 from 3 technical replicates each. **H**, Phosphorylation of Smad2 in wild-type, *sparc*^+/-^, and *des*^+/-^ CSCs and CBs after addition of glycosylated SPARC for 3 hours and 3 days, respectively, and in SPARC overexpressing *sparc^+/+^S^ect^*CSCs. **I-L**, Expression of cardiomyogenic transcription factor genes and *desmin* as control in 8.5 days old CBs by RT-qPCR. Data from 2 *des^+/-^* cell lines and 2 experiments with 2 and 3 biological replicates, respectively, and each 3 replicates. *, Student′s t-test p value p<0.05; ***, Student′s t-test p value p<0.001.

Comparing the expression of myocardial transcription factor genes in control and SPARC-treated *des^+/-^* CBs shows that glycosylated but not de-glycosylated SPARC promoted the expression of *gata4*, *mef2C* (**Figure 7I-J**) and to a lesser extent *nkx2.5* at day 8.5 of cardiomyogenesis (**Figure 7K**) and, as already shown in Figure 5C, that *desmin* expression was strongly increased (**Figure 7L**). This suggests that glycosylated and secreted SPARC increases and accelerates cardiomyogenesis in CBs by promoting EMT, increasing the phosphorylation of Smad2, and the expression of myocardial transcription factor genes. In conclusion, SPARC ameliorated *desmin* haploinsufficiency in CBs in a glycosylation-dependent manner, and it is the extracellular rather than the intracellular SPARC that promotes cardiomyogenesis in CSCs.

## Discussion

Here we investigated the role of the muscle cell specific type III intermediate filament protein desmin during cardiomyogenesis in cardiac stem cells (CSCs). Based on the evidence that *desmin* overexpression in embryoid bodies (EBs) ^15^ promoted cardiomyogenesis and the expression of *brachyury* and *nkx2.5* ^16^ in a cell autonomous manner, and the observation that promotion of cardiomyogenesis was also influenced by the density of EBs in the culture plates, we hypothesized that *desmin* expression may also contribute to cardiomyogenesis via an autocrine or paracrine non-cell autonomous mechanism. Testing this hypothesis by co-culture of *desmin*-overexpressing EBs with wild-type and *desmin*-null EBs showed that desmin induces the expression and secretion of the ECM-associated matri-cellular protein SPARC and that secreted SPARC promoted cardiomyogenesis in a Smad2-dependent manner (Figure 1). The main limitation of this approach was that EBs are composed of cells from all three germ layers, all of which ectopically express desmin. This made it impossible to distinguish between a general paracrine and an autocrine, cell autonomous promotion of cardiomyogenesis by SPARC. Therefore, we changed our model system to CSCs, which already express substantial amounts of desmin in the undifferentiated, self-renewing state and exclusively give rise to cardiac progenitor cells that differentiate into cardiomyocytes, cardiac endothelial cells and cardiac smooth muscle cells ^12^. Even more importantly, in vivo data suggests that desmin deficiency impairs the cardiomyogenic commitment of cardiac stem cell populations ^53^. Self-renewing CSCs indeed expressed substantial amounts of SPARC, which was localized in the Golgi apparatus, cytoplasm and nucleus of CSCs. SPARC was also secreted, bound to the ECM and later taken up by differentiating CSCs (Figure 2). Strongly SPARC-expressing cells were also found sparsely localized at the border of muscle fibers in mouse hearts, suggesting that our model CSCs may be derived from these cell populations or at least in this regard mimic in vivo CSCs correctly. These findings led us to ask whether autocrine secretion of SPARC from CSCs also promotes cardiomyogenesis in CSCs. Paracrine and autocrine SPARC promoted cardiomyogenesis in differentiating CSCs when aggregated into cardiac bodies (CBs) at the very beginning of cardiomyogenesis in a dose-dependent manner (Figure 3). Constitutive overexpression of *sparc* also increased the longevity and robustness of rhythmic contraction of cardiomyocytes, consistent with the previously shown positive ionotropic influence of SPARC on cardiomyocytes ^49^. Notably, *sparc* haploinsufficiency in CSCs could be partially rescued by paracrine addition of SPARC, demonstrating that it mainly acts as an extracellular signaling protein, but also strongly suggesting that there are additional cell-autonomous functions of SPARC that could not be compensated by paracrine application of the protein. Not being able to distinguish between the possibly different functions of SPARC taken up by the cells and cytoplasmic SPARC is another limitation of this study. Incomplete rescue by paracrine SPARC, together with the repeated failure to generate SPARC-null CSCs, may point to another unknown and perhaps essential role of SPARC in CSCs.

Paracrine SPARC also increased the expression of early myocardial transcription factor genes in self-renewing CSCs, and unexpectedly also increased the expression of a *desmin*-reporter gene, at least in a small subset of CSCs (Figure 4). Analysis of the entire population of CSCs by RT-qPCR showed a rapid increase in mesodermal *brachyury* (*T*) mRNA expression and myocardial *mesp1*, *nkx2.5* and *gata4* transcription factor expression. However, FACS-based single cell analysis of reporter cell lines showed expression of *T*, *mesp1*, *nkx2.5* and *desmin* only in a variable subset of CSCs at a given time point. This phenomenon may either be due to the fact that the *T*, *mesp1* and *desmin* reporter constructs contained only parts of the promoter and enhancer regions of their genes or, more likely, that CSCs, similar to ESCs ^54^, exhibit stochastic gene expression patterns that are also influenced by their cellular circadian clock ^55^. In addition, it became clear that *nkx2.5*, whose expression had been shown to be influenced by both SPARC and desmin in ESC-derived cardiac progenitors and primary cardiomyocytes ^14, 16^, was only transiently induced by paracrine SPARC in CSCs. These discrepancies are a clear limitation of this study and could only be resolved in the future by a large single cell transcriptomic study at several consecutive developmental stages of differentiating CSCs.

The finding that paracrine SPARC also promotes *desmin* expression in CSCs led us to hypothesize that desmin and SPARC may form a modifiable and reinforcing positive feedback loop in CSCs that may contribute to CSC-based cardiomyogenesis in vitro and perhaps in the adult heart. Analysis of mRNA expression and protein synthesis in CSCs with one, two or three alleles of *sparc* and *desmin*, respectively, showed that desmin mRNA and protein levels were directly correlated with SPARC mRNA and protein levels, and conversely, that SPARC mRNA and protein levels, as well as its secretion into the medium, were directly correlated with desmin mRNA and protein levels (Figure 5). These results strongly suggest a genetic interaction of these two genes and a positive feedback loop mediated by their proteins. In vitro differentiation of CSCs with mono-allelic expression of *desmin* revealed a *desmin*-haplo-insufficient cardiomyogenic phenotype in CBs (Figure 6), which was very similar to that of CBs with mono-allelic expression of *sparc*. In contrast to the EB model, overexpression of *desmin* in CBs resulted in a dominant-negative cardiomyogenic phenotype, leading to arrhythmic contraction and premature death of cardiomyocytes, most likely caused by desmin protein aggregation, which is also observed in human desminopathies ^56–58^.

The *desmin* haploinsufficiency in CBs together with reduced SPARC secretion from these cells suggested that the lack of sufficient amounts of autocrine SPARC may be the cause of reduced cardiomyogenesis. Paracrine glycosylated, but not de-glycosylated, SPARC did indeed restore cardiomyogenesis and contraction frequency of cardiomyocytes (Figure 7), and additionally caused increased phosphorylation of Smad2 and significant upregulation of *gata4*, *mef2C*, *nkx2.5* and *desmin* during cardiomyogenesis in CBs. This strongly suggests that desmin is, at least in part, responsible for the expression and secretion of glycosylated SPARC in CSCs and that autocrine and paracrine SPARC in turn promotes cardiomyogenesis directly via modulation of the Tgf-ß-Smad2 signal transduction in CSCs and increases rhythmic contraction and longevity of cardiomyocytes. However, it remains open if SPARC acts indirectly via modifying the ECM or directly via signal transduction and/or endocytotic uptake by the cells.

In conclusion, both genes and proteins together appear to form a positive feedback loop in CSCs, which promotes cardiomyogenesis, and that CSCs may have a gland like function in the heart by secreting SPARC. Further, these in vitro data, together with the fact that *desmin* mutations and aberrant SPARC levels contribute to cardiac diseases, suggest that the genetic link and interplay between desmin and SPARC may also contribute to the maintenance of homeostasis in the adult heart in vivo, and may thus be exploited for induced myocardial regeneration or as targets in future therapies.

## Nonstandard Abbreviations and Acronyms

CSCs: cardiac stem cells
CBs: cardiac bodies
ESCs: embryonic stem cells
EBs: embryoid bodies
EMT: epithelial to mesenchymal transition
SPARC: secreted protein acidic and rich in cysteine

## Acknowledgments

We would like to thank Wei Chiang Liao, Theresa Matzinger, Petra Kalmann, Sabrina Beil, Brigitte Gundacker, Thomas Sauer, and Irmgard Fischer for technical help, was well as the students Marina Leiwe, Lisa Marie Wittmann, Melissa Sperlich, Albert Dorca Fabrega, Dominik Förger, Theresa Hillinger, Cornelia Kodek, Chiara Rossi, Sahrah Linauer, Daniel Neubauer, Jakob Girlinger, Roxana Rehak, Melanie Hobik, Nina Koppensteiner, Barbara Nimeth, Leila Djerlak, and Nathalie Tichy Prada for helping with the experimental work of the coauthors.

## Author contribution

L.L., M.S., F.H., C.R., V.K., K.F., M-T.K., J.H., M.G., J.K., E.A., T.G., D.W. performed the experiments and analyzed the data, F.H. and G.W. rote the manuscript and G.W. conceived and designed the project and overlooked all data.

## Additional Information

Supplemental Information are

Supplemental Figure 1 + Legend

Major Resources Table

## Supplemental Information

**Supplemental Figure 1.**
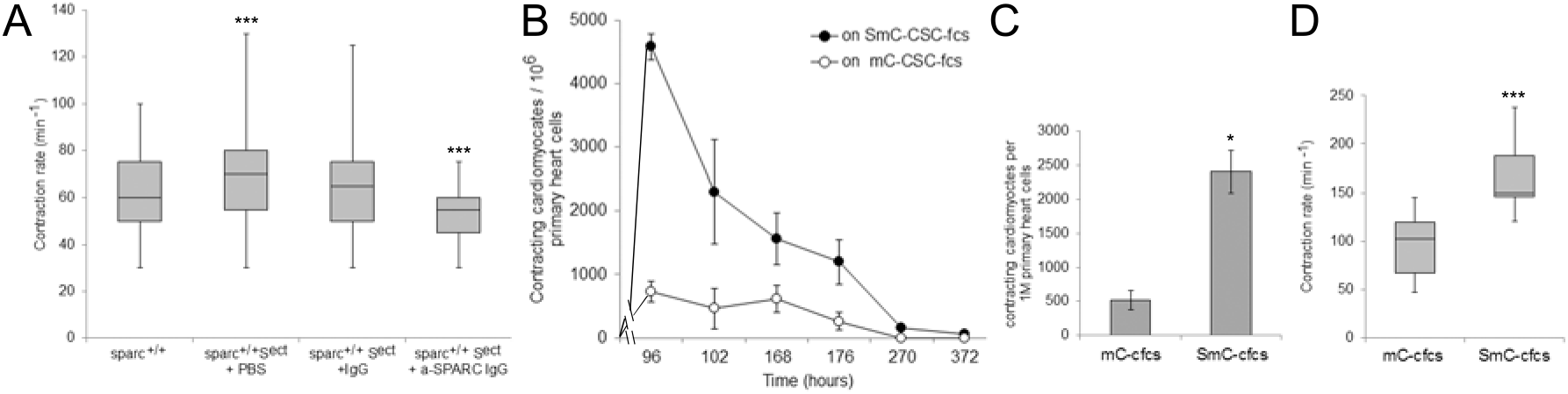
SPARC increases the frequency of cardiomyocyte contraction and fosters the survival and contraction of primary cardiomyocytes. (**A**) Frequency of rhythmic contraction of SPARC overexpressing *sparc^+/+^S^ect^* cardiomyocytes compared to wild-type *sparc^+/+^*cardiomyocytes and *sparc^+/+^S^ect^* cardiomyocyte in the presence of anti-SPARC antibodies. Box plots show the median and the second and third quartiles; whiskers indicate maximum and minimum values, respectively. Data were collected between days 11 and 26 of CB development, each N≥60; n=3. ***, Student′s t-test p value p<0.001.(**B**) Time course of cardiomyocyte survival after plating on CSC derived feeder cells either overexpressing SPARC:mCherry fusion protein (SmC-CSCfcs) or only mCherry protein (mC-CSCfcs) and (**C**) Number of contracting cardiomyocytes isolated from 9 N5 mice and plated on CSC-derived feeder cells expressing either only mCherry or SPARC-mCherry fusion protein. N=4, Error bars, standard deviation.*, Student′s t-test p value p<0.05. (**D**) Frequency of rhythmic contraction of primary cardiomyocytes in the presence of mC-CSCfcs and SmC-CSCfcs. Condition as in (**C**), ***, Student′s t-test p value p<0.001

## Major Resources Table

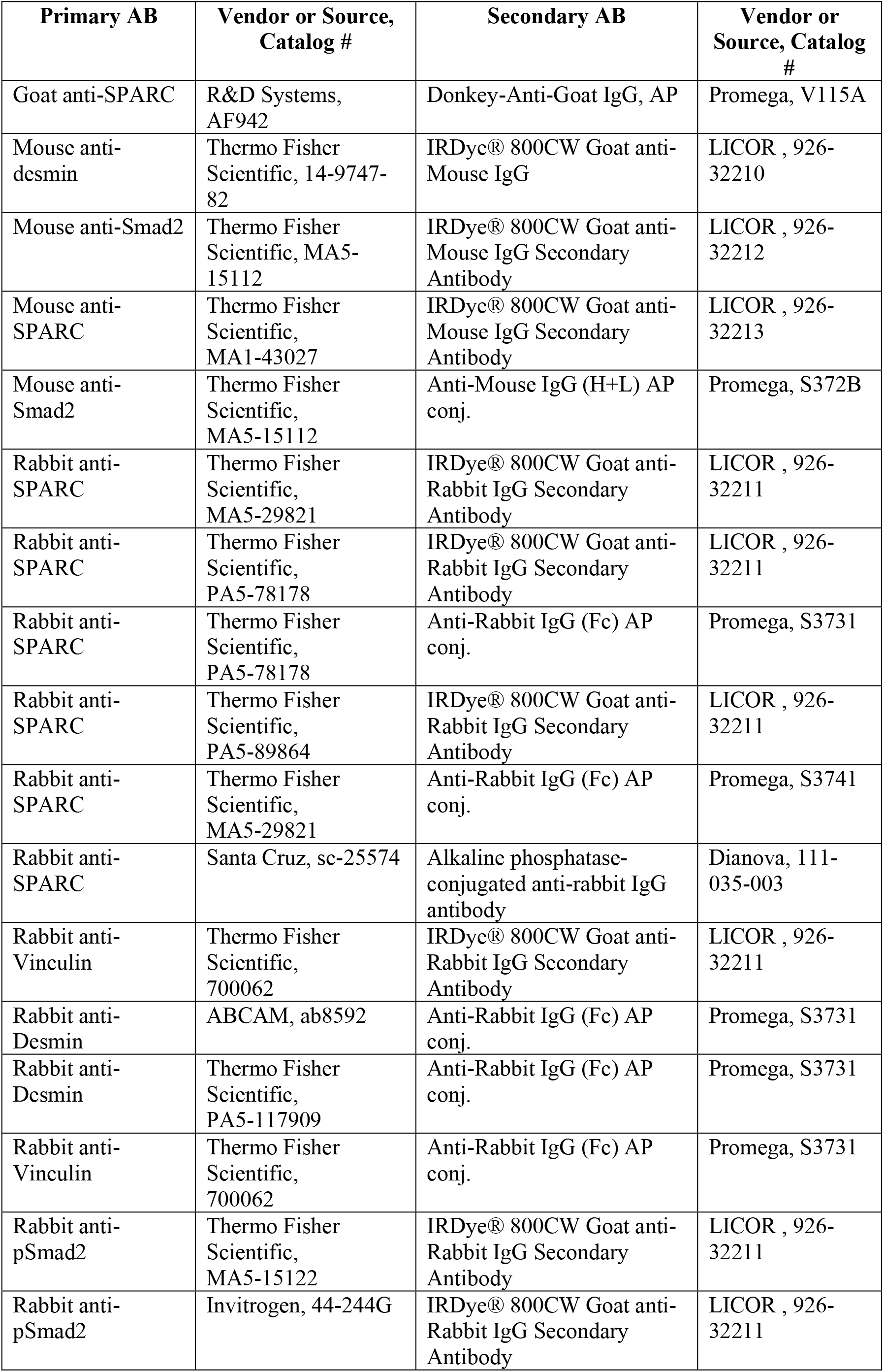
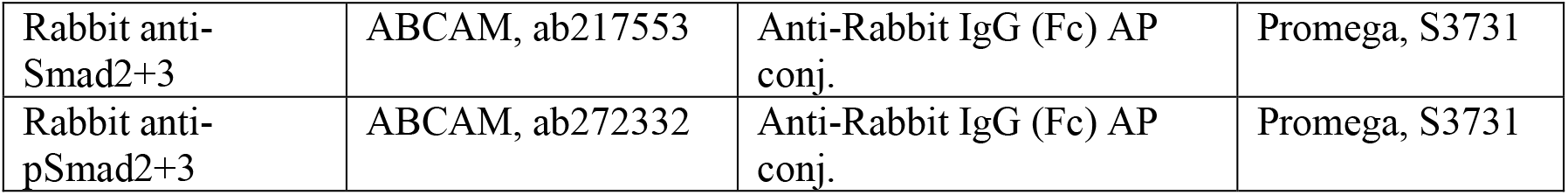
Antibodies.

Antibodies
| Primary AB | Vendor or Source, Catalog # | Secondary AB | Vendor or Source, Catalog # |
| --- | --- | --- | --- |
| Goat anti-SPARC | R&D Systems, AF942 | Donkey-Anti-Goat IgG, AP | Promega, V115A |
| Mouse anti-desmin | Thermo Fisher Scientific, 14-9747-82 | IRDye® 800CW Goat anti-Mouse IgG | LICOR , 926-32210 |
| Mouse anti-Smad2 | Thermo Fisher Scientific, MA5-15112 | IRDye® 800CW Goat anti-Mouse IgG Secondary Antibody | LICOR , 926-32212 |
| Mouse anti-SPARC | Thermo Fisher Scientific, MA1-43027 | IRDye® 800CW Goat anti-Mouse IgG Secondary Antibody | LICOR , 926-32213 |
| Mouse anti-Smad2 | Thermo Fisher Scientific, MA5-15112 | Anti-Mouse IgG (H+L) AP conj. | Promega, S372B |
| Rabbit anti-SPARC | Thermo Fisher Scientific, MA5-29821 | IRDye® 800CW Goat anti-Rabbit IgG Secondary Antibody | LICOR , 926-32211 |
| Rabbit anti-SPARC | Thermo Fisher Scientific, PA5-78178 | IRDye® 800CW Goat anti-Rabbit IgG Secondary Antibody | LICOR , 926-32211 |
| Rabbit anti-SPARC | Thermo Fisher Scientific, PA5-78178 | Anti-Rabbit IgG (Fc) AP conj. | Promega, S3731 |
| Rabbit anti-SPARC | Thermo Fisher Scientific, PA5-89864 | IRDye® 800CW Goat anti-Rabbit IgG Secondary Antibody | LICOR , 926-32211 |
| Rabbit anti-SPARC | Thermo Fisher Scientific, MA5-29821 | Anti-Rabbit IgG (Fc) AP conj. | Promega, S3741 |
| Rabbit anti-SPARC | Santa Cruz, sc-25574 | Alkaline phosphatase-conjugated anti-rabbit IgG antibody | Dianova, 111-035-003 |
| Rabbit anti-Vinculin | Thermo Fisher Scientific, 700062 | IRDye® 800CW Goat anti-Rabbit IgG Secondary Antibody | LICOR , 926-32211 |
| Rabbit anti-Desmin | ABCAM, ab8592 | Anti-Rabbit IgG (Fc) AP conj. | Promega, S3731 |
| Rabbit anti-Desmin | Thermo Fisher Scientific, PA5-117909 | Anti-Rabbit IgG (Fc) AP conj. | Promega, S3731 |
| Rabbit anti-Vinculin | Thermo Fisher Scientific, 700062 | Anti-Rabbit IgG (Fc) AP conj. | Promega, S3731 |
| Rabbit anti-pSmad2 | Thermo Fisher Scientific, MA5-15122 | IRDye® 800CW Goat anti-Rabbit IgG Secondary Antibody | LICOR , 926-32211 |
| Rabbit anti-pSmad2 | Invitrogen, 44-244G | IRDye® 800CW Goat anti-Rabbit IgG Secondary Antibody | LICOR , 926-32211 |
| Rabbit anti-Smad2+3 | ABCAM, ab217553 | Anti-Rabbit IgG (Fc) AP conj. | Promega, S3731 |
| Rabbit anti-pSmad2+3 | ABCAM, ab272332 | Anti-Rabbit IgG (Fc) AP conj. | Promega, S3731 |

Plasmids DNA/cDNA Clones.

Plasmids DNA/cDNA Clones
| Name | Sequence | Source / Repository |
| --- | --- | --- |
| pNKE24 |  | <sup>44</sup> |
| pGL3-Basic | Addgene | Promega, E1751 |
| phRL-TK | Addgene | Promega, E2241 |
| mouse 1.2kb pT-BraP-Puro-IRES2-eGFP |  | <sup>40</sup> |
| human 3.4 kb pMesP1-eGFP |  | <sup>41</sup> |
| pCSX-EGFP-PPDT |  | <sup>42</sup> |
| pCMV3-SPARC | NM_009242.4 | SinoBiological, MG50494-UT |
| desmin expression cassette | GenBank: Z18892.2 | <sup>15</sup> |
| pmCherry N1 vector | Addgene | Clontech, 632523 |

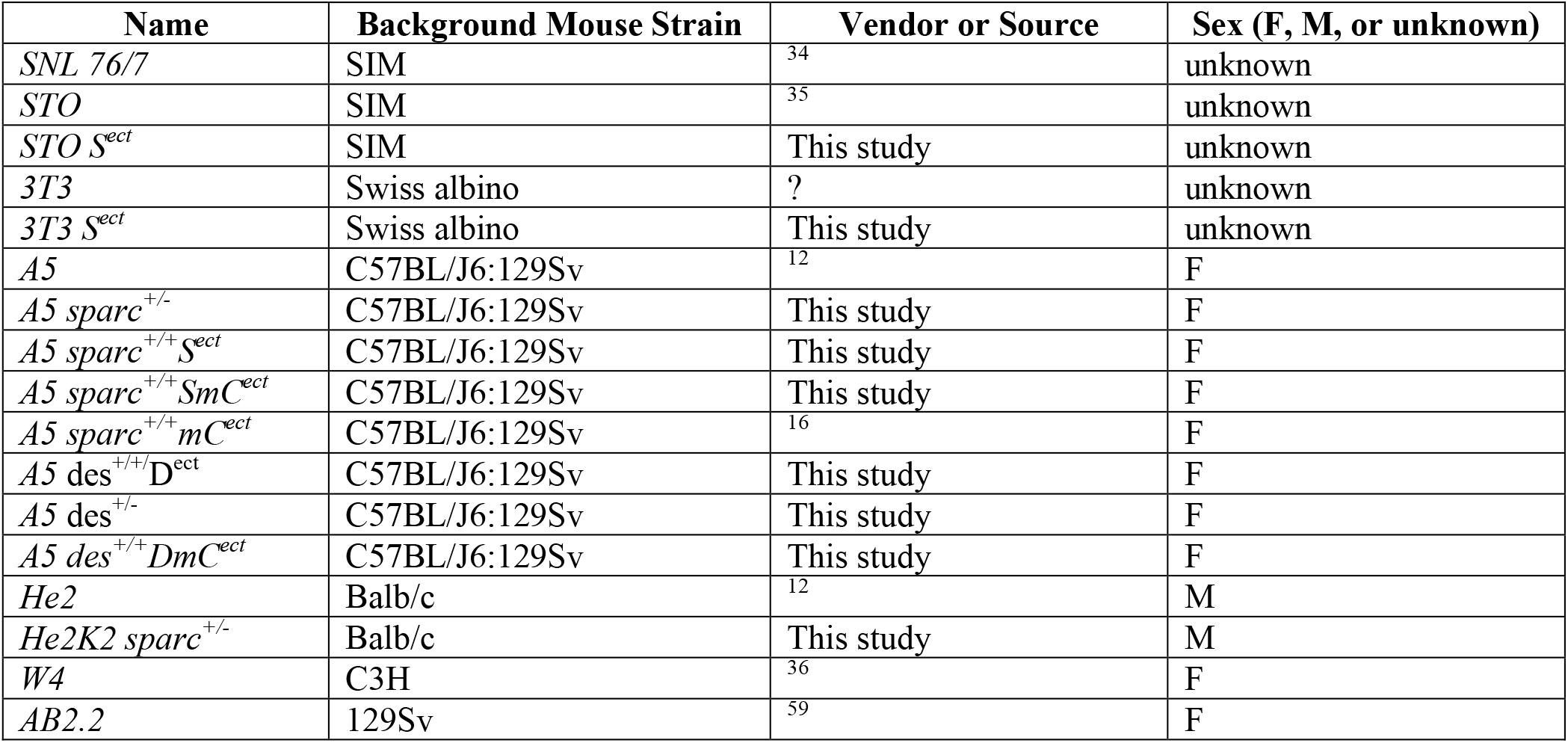
Cultured Cells.

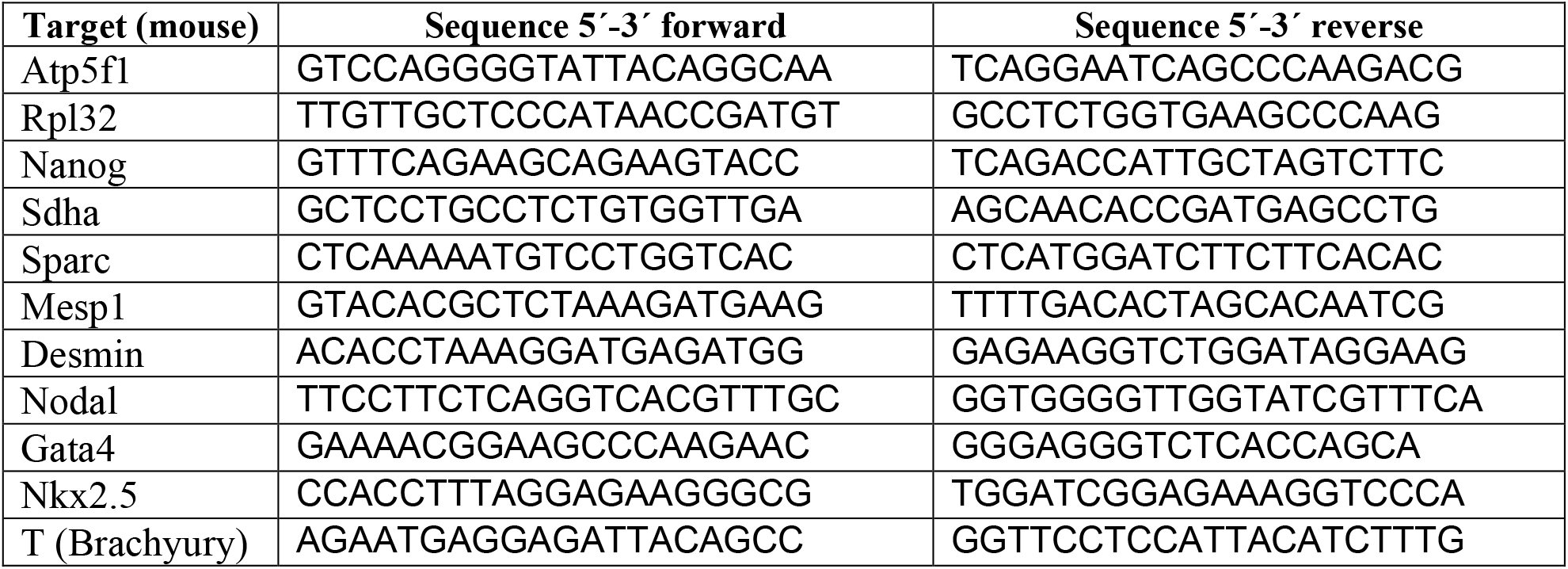
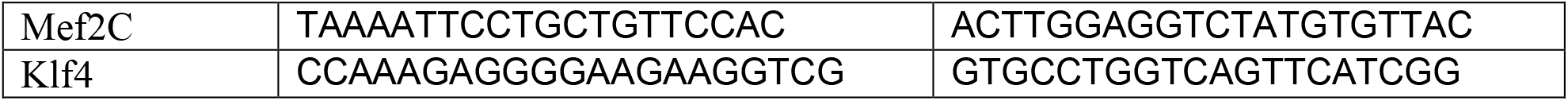
qPCR Primer.

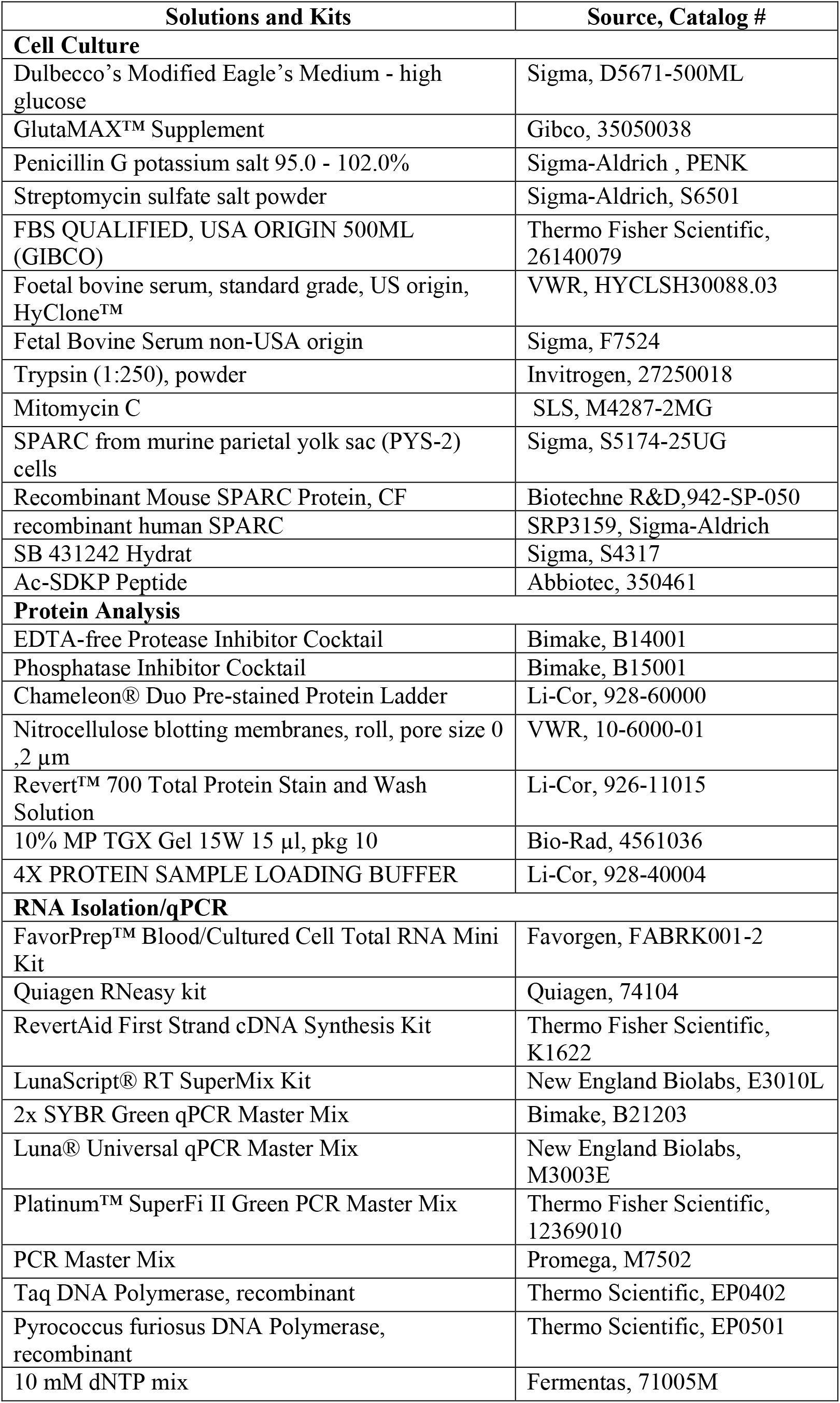
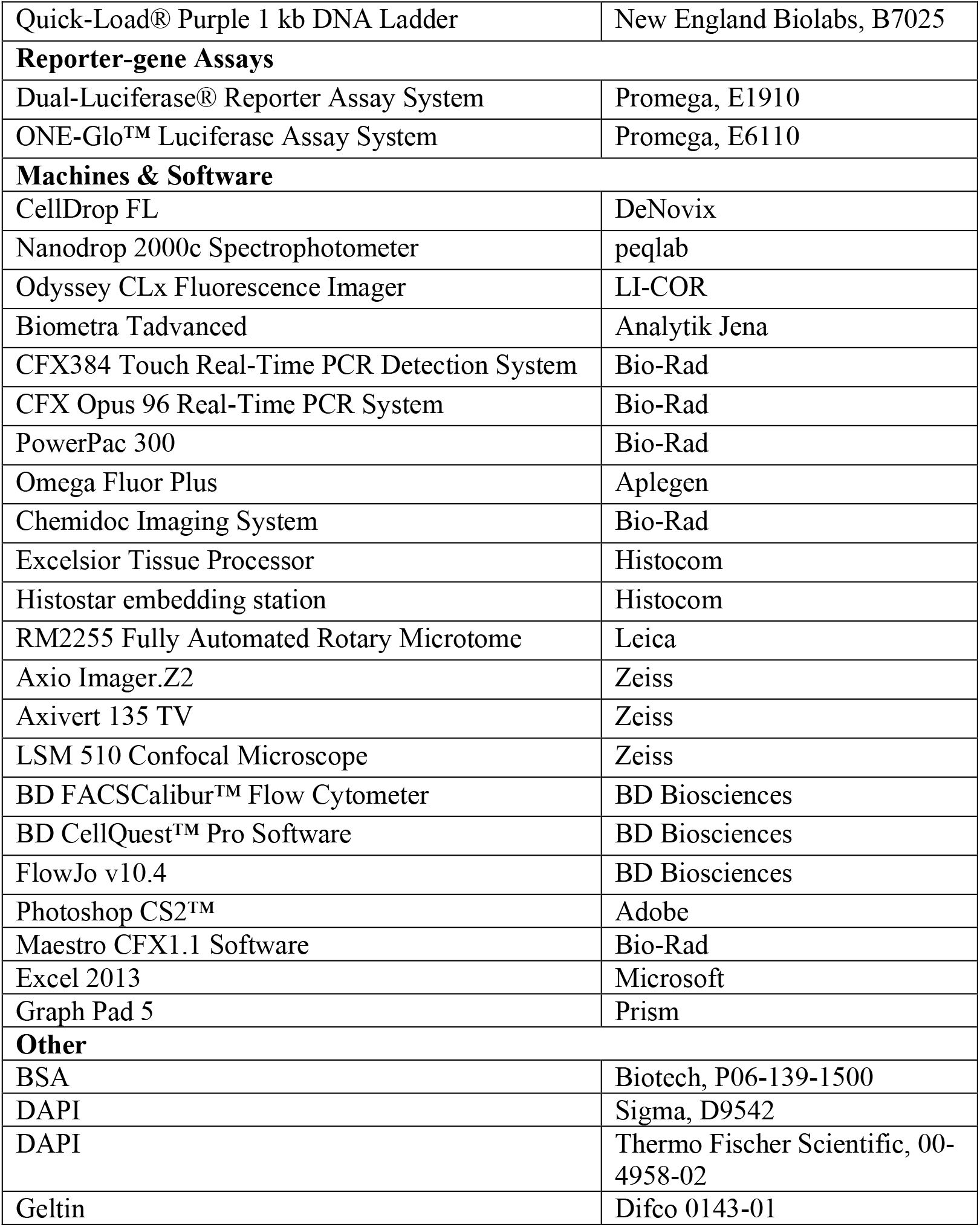
Other.

Other
| Solutions and Kits | Source, Catalog # |
| --- | --- |
| **Cell Culture** |  |
| Dulbecco's Modified Eagle's Medium - high glucose | Sigma, D5671-500ML |
| GlutaMAX™ Supplement | Gibco, 35050038 |
| Penicillin G potassium salt 95.0 - 102.0% | Sigma-Aldrich , PENK |
| Streptomycin sulfate salt powder | Sigma-Aldrich, S6501 |
| FBS QUALIFIED, USA ORIGIN 500ML (GIBCO) | Thermo Fisher Scientific, 26140079 |
| Foetal bovine serum, standard grade, US origin, HyClone™ | VWR, HYCLSH30088.03 |
| Fetal Bovine Serum non-USA origin | Sigma, F7524 |
| Trypsin (1:250), powder | Invitrogen, 27250018 |
| Mitomycin C | SLS, M4287-2MG |
| SPARC from murine parietal yolk sac (PYS-2) cells | Sigma, S5174-25UG |
| Recombinant Mouse SPARC Protein, CF | Biotechne R&D,942-SP-050 |
| recombinant human SPARC | SRP3159, Sigma-Aldrich |
| SB 431242 Hydrat | Sigma, S4317 |
| Ac-SDKP Peptide | Abbiotec, 350461 |
| **Protein Analysis** |  |
| EDTA-free Protease Inhibitor Cocktail | Bimake, B14001 |
| Phosphatase Inhibitor Cocktail | Bimake, B15001 |
| Chameleon® Duo Pre-stained Protein Ladder | Li-Cor, 928-60000 |
| Nitrocellulose blotting membranes, roll, pore size 0.2 µm | VWR, 10-6000-01 |
| Revert™ 700 Total Protein Stain and Wash Solution | Li-Cor, 926-11015 |
| 10% MP TGX Gel 15W 15 µl, pkg 10 | Bio-Rad, 4561036 |
| 4X PROTEIN SAMPLE LOADING BUFFER | Li-Cor, 928-40004 |
| **RNA Isolation/qPCR** |  |
| FavorPrep™ Blood/Cultured Cell Total RNA Mini Kit | Favorgen, FABRK001-2 |
| Quiagen RNeasy kit | Quiagen, 74104 |
| RevertAid First Strand cDNA Synthesis Kit | Thermo Fisher Scientific, K1622 |
| LunaScript® RT SuperMix Kit | New England Biolabs, E3010L |
| 2x SYBR Green qPCR Master Mix | Bimake, B21203 |
| Luna® Universal qPCR Master Mix | New England Biolabs, M3003E |
| Platinum™ SuperFi II Green PCR Master Mix | Thermo Fisher Scientific, 12369010 |
| PCR Master Mix | Promega, M7502 |
| Taq DNA Polymerase, recombinant | Thermo Scientific, EP0402 |
| Pyrococcus furiosus DNA Polymerase, recombinant | Thermo Scientific, EP0501 |
| 10 mM dNTP mix | Fermentas, 71005M |
| Quick-Load® Purple 1 kb DNA Ladder | New England Biolabs, B7025 |
| **Reporter-gene Assays** |  |
| Dual-Luciferase® Reporter Assay System | Promega, E1910 |
| ONE-Glo™ Luciferase Assay System | Promega, E6110 |
| **Machines & Software** |  |
| CellDrop FL | DeNovix |
| Nanodrop 2000c Spectrophotometer | peqlab |
| Odyssey CLx Fluorescence Imager | LI-COR |
| Biometra Tadvanced | Analytik Jena |
| CFX384 Touch Real-Time PCR Detection System | Bio-Rad |
| CFX Opus 96 Real-Time PCR System | Bio-Rad |
| PowerPac 300 | Bio-Rad |
| Omega Fluor Plus | Aplegen |
| Chemidoc Imaging System | Bio-Rad |
| Excelsior Tissue Processor | Histocom |
| Histostar embedding station | Histocom |
| RM2255 Fully Automated Rotary Microtome | Leica |
| Axio Imager.Z2 | Zeiss |
| Axivert 135 TV | Zeiss |
| LSM 510 Confocal Microscope | Zeiss |
| BD FACSCalibur™ Flow Cytometer | BD Biosciences |
| BD CellQuest™ Pro Software | BD Biosciences |
| FlowJo v10.4 | BD Biosciences |
| Photoshop CS2™ | Adobe |
| Maestro CFX1.1 Software | Bio-Rad |
| Excel 2013 | Microsoft |
| Graph Pad 5 | Prism |
| **Other** |  |
| BSA | Biotech, P06-139-1500 |
| DAPI | Sigma, D9542 |
| DAPI | Thermo Fischer Scientific, 00-4958-02 |
| Geltin | Difco 0143-01 |

## Novelty and Significance

### What Is Known?

- Although preserved throughout mammalian evolution, cardiac stem cells do not contribute significantly to the repair of the injured heart.
- SPARC and desmin are expressed at a very early stage of cardiogenesis and contribute to myocardial gene expression and cardiomyogenesis in vitro.
- Mutations in both genes negatively affect cardiac repair in a similar manner.
- Desmin is expressed in cardiac stem cells and promotes cardiomyogenesis via activation of *nkx2.5* expression in a cell autonomous manner.

### What New Information Does This Article Contribute?

- Desmin induces SPARC expression and promotes cardiomyogenesis in embryoid bodies also in a non-cell autonomous manner.
- Cardiac stem cells secrete SPARC which binds to the extracellular matrix and is taken up by differentiating cardiac progenitor cells.
- Paracrine and autocrine SPARC promotes cardiomyogenesis in cardiac stem cells in a concentration-dependent manner and increases the phosphorylation of Smad2.
- SPARC influences the expression of *desmin* and *brachyury*, *mesp1*, *gata4*, and *nkx2.5* in self-renewing and differentiating cardiac stem cells.
- SPARC and desmin reciprocally promote their expression and protein synthesis and the secretion of SPARC but excess of paracrine SPARC inhibits *sparc* expression.
- Reduced expression of desmin inhibits cardiomyogenesis and SPARC rescues desmin-related cardiomyogenic haploinsufficiency in a glycosylation-dependent manner by increasing the pSmad2/Smad2 ratio and activating the expression of myocardial transcription factor genes.

These data suggest that

- Both desmin and SPARC synergistically promote cardiomyogenesis in cardiac stem cells.
- They establish a regulatory genetic network including a positive desmin-SPARC and a negative SPARC-*sparc* loop regulating the expression of myocardial transcription factor genes such as *gata4*, *nkx2.5*, and *mef2c*.
- SPARC-secreting cardiac stem cells may have a gland-like function in the adult myocardium helping to regain myocardial homeostasis in the injured or aging heart.

